# Network-based anomaly detection algorithm reveals proteins with major roles in human tissues

**DOI:** 10.1101/2023.12.19.572354

**Authors:** Dima Kagan, Juman Jubran, Esti Yeger-Lotem, Michael Fire

**Affiliations:** Department of Software and Information Systems Engineering, Ben-Gurion University of the Negev, Beer Sheva 84105, Israel; Department of Clinical Biochemistry and Pharmacology, Ben-Gurion University of the Negev, Beer Sheva 84105, Israel; The National Institute for Biotechnology in the Negev, Ben-Gurion University of the Negev, Beer Sheva 84105, Israel

## Abstract

**Background:** Anomaly detection in graphs is critical in various domains, notably in medicine and biology, where anomalies often encapsulate pivotal information. Here, we focused on network analysis of molecular interactions between proteins, which is commonly used to study and infer the impact of proteins on health and disease. In such a network, an anomalous protein might indicate its impact on the organism’s health.

**Results:** We propose Weighted Graph Anomalous Node Detection (WGAND), a novel machine learning-based method for detecting anomalies in weighted graphs. WGAND is based on the observation that edge patterns of anomalous nodes tend to deviate significantly from expected patterns. We quantified these deviations to generate features, and utilized the resulting features to model the anomaly of nodes, resulting in node anomaly scores. We created four variants of the WGAND methods and compared them to two previously-published (baseline) methods. We evaluated WGAND on data of protein interactions in 17 human tissues, where anomalous nodes corresponded to proteins with major roles in tissue contexts. In 13 of the tissues, WGAND obtained higher AUC and P@K than baseline methods. We demonstrate that WGAND effectively identified proteins that participate in tissue-specific processes and diseases.

**Conclusion:** We present WGAND, a new approach to anomaly detection in weighted graphs. Our results underscore its capability to highlight critical proteins within protein-protein interaction networks. WGAND holds the promise to enhance our understanding of intricate biological processes and might pave the way for novel therapeutic strategies targeting tissue-specific diseases. Its versatility ensures its applicability across diverse weighted graphs, making it a robust tool for detecting anomalous nodes.

## 1 Introduction

The detection of anomalies is a crucial problem in diverse domains, ranging from cybersecurity to social network analysis. Anomalies refer to elements or objects “that appear to deviate markedly from other members of the sample in which it occurs” [1]. The study of anomalies has fascinated scientists for centuries, as they often hold unique insights into the dynamics of complex systems [2]. Anomaly detection is an invaluable tool for various disciplines, from aviation to medicine, to uncover insights that are not easily found through traditional methods [2]. For instance, in aviation, an anomaly found in airplane sensors may indicate a fault. In medicine, an anomaly can be considered as a patient with a rare disease [3].

In recent years, anomaly detection in graphs has gained increasing attention due to its applicability in various domains such as cybersecurity, social networks, and biological networks [3]. The structure and interconnected nature of graphs can provide rich information for anomaly detection. However, the graph’s complex nature, including its size, diversity, and noise, makes this task challenging [3].

Over the years, many studies have addressed anomaly detection, especially in graphs. However, only a few studies focused on anomaly detection in weighted graphs [4]. For example, Akoglu et al. [5] presented OddBall, an algorithm that detects anomalous nodes in weighted graphs based on generating features that transform into a score to pinpoint outliers. The features are generated from the neighborhood of each node. The following year, Davis et al. [6] presented Yagada, an algorithm that searches for structural and numerical anomalies. Another study by Lee et al. [4] also presented a method for detecting outliers in weighted graphs, but their method focuses only on directed weighted graphs. The algorithm iterates through the graph, searching for the “best” substructures, which generate the largest compression when replaced with a super-node. The anomaly score is generated based on how well it was compressed.

In this study, we propose Weighted Graph Anomalous Node Detection (WGAND), a novel generic anomaly detection machine learning-based algorithm for weighted graphs. The method assumes that on average edge weights of anomalous nodes should deviate from their expected norm. In other words, if we can create a model that estimates expected edge weights and compare them with the observed value, we can quantify the deviation of edge weights from the norm (see Fig. 1). Then, by accumulating the deviation, we can detect anomalous nodes. To this end, we trained a model that estimates the expected edge weight for all edges in the graph. Using the difference between the actual and expected weight, we generate eight features that are used to generate an anomaly score for all the nodes in the graph.

**Figure 1:**
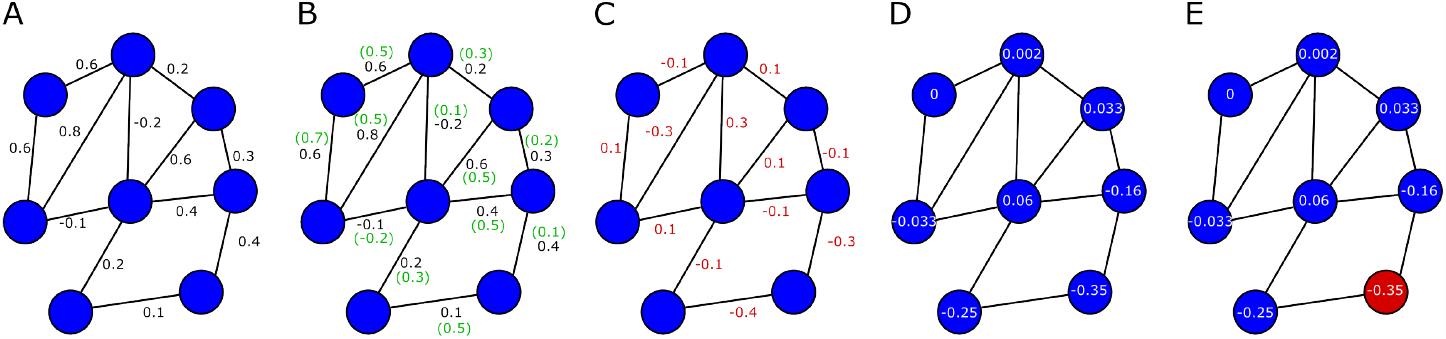
Schematic illustration of detecting anomalous nodes in weighted graphs. **A)** Given a weighted graph, a weight estimator is trained. **B)** The predicted estimated weights of the edges (in green) are calculated based on the weight estimator. **C)** Each edge is assigned with the difference value between the estimated and actual weights. **D)** For each node, the difference value of its edges is aggregated and used to create an anomaly detection score. Practically, we use a set of models to quantify a set of such features to a single value that is used as the score. **E)** Nodes with values higher or lower than a specific threshold are considered anomalous. For instance, the red node is the node with the lowest value in the graph.

To demonstrate and evaluate the value of WGAND, we turned to weighted protein-protein interaction (PPI) networks. Proteins are the prime molecules in living cells and conduct their activities through physical interactions with other molecules. PPI networks, whereby proteins are represented by nodes and their interactions by edges, provide a useful framework to study protein functions and cellular information transfer [7]. Weighted PPI networks, where edge weight reflects the likelihood of an interaction, offer a refined view of context-specific interactions [8]. For example, weighted PPI networks representing different human tissues helped understand tissue-specific cellular processes and functions in health and diseases [9, 10, 11]. Here, we hypothesized that proteins involved in major biological roles within a specific tissue will appear as anomalous nodes in the PPI network that represent the inflicted tissue. Among these proteins are those involved in tissue-specific diseases, such as BRCA1 whose aberrations increase the risk for breast and ovarian cancers, and those that participate in main tissue processes, such as neuron signaling in the brain and spermatogenesis in the testis. To test our hypothesis, we applied the WGAND algorithm to weighted PPI networks of 17 different human tissues and to 1,541 proteins involved in tissue-specific diseases. We found that WGAND obtained higher Area Under the ROC curve (AUC) and precision at k (P@K), than other methods when applied to identify disease-related proteins in most tissues. High-ranking anomalous proteins were more likely to be associated with diseases than other proteins, were enriched for proteins involved in tissue-specific processes, and tended to be involved in processes that were active preferentially in the designated tissue. Whereas our study used weighted PPI networks as a case study, WGAND is a generic method that could be applied to any weighted graph.

The key contributions presented in this paper are three-fold:

- We present a set of novel anomaly detection scores for anomaly detection in weighted graphs. This study is one of the few studies to use ground truth anomalies to examine graph anomaly detection instead of injecting simulated anomalies. The algorithm was more accurate and precise than previous methods.
- Application of graph-based anomaly detection to tissue-specific networks revealed proteins with major tissue-specific roles. To the best of our knowledge, this is the first such usage of an anomaly-detection algorithm.
- We present an open Python library implementing WGAND and release all data used in this research to help advance additional research on this vital task.

The remainder of this paper is organized as follows. In Section 2, we describe the datasets, methods, algorithms, and experiments used throughout this study. In Section 3, we present our results. Lastly, in Section 4, we discuss the obtained results and offer future research directions.

## 2 Methods

### 2.1 Problem definition

It is difficult to define precisely what constitutes an anomaly [12]. Definitions of anomalies may vary across domains and disciplines. However, anomalies are generally considered to be unusual observations that differ from the general population to the degree that suggests a different mechanism may have created them [5]. In this study, we treated an anomaly as a vertex that behaves differently from most of the vertices population. Formally, we defined the problem as follows: given a weighted graph *G* = (*V, E, W*) consisting of a set of vertices *V*, a set of edges *E*, and a set of edge weights *W*, each node *v* ∈ *V* has a set of edges (*v, u*) ∈ *E* which has weight *W*_*u,v*_ ∈ ℝ. A node *v ∈ V* is considered anomalous if a function *f* (*v, G*) which represents a node anomaly score is larger than a predefined *threshold*.

### 2.2 General Framework

In this section, we describe the general framework of our anomaly detection method, which is based on the idea that anomalous nodes should interact differently with their neighbors.

To formalize this idea, let us consider a weighted graph *G* = (*V, E, W*), where *V* is the set of vertices, *E* is the set of edges, and *W* is the set of weights. Given a vertex *v*, we define its neighborhood Γ(*v*) = { (*v, u*) | (*v, u*) ∈ *E* } as the set of vertices that are connected to *v* by an edge, and their weights as *W*_*v*_ = {*W*_*v,u*_ | *u* ∈ Γ(*v*) }. Where *W*_*a*_ and *W*_*n*_ represent the edge weights between a suspected anomalous node a with its neighbors and a normal node *n* with its neighbors, respectively.

Inspired by the work of Kagan et al. [13], we hypothesized that *W*_*a*_ should deviate from *W*_*n*_ if *a* is truly anomalous. In other words, the average edge weights for *a* should differ from those of *n*. To test this hypothesis, we assumed that a general predictive function *P*_*f*_ (*v, u*) exists, in which it can estimate the weight of an edge (*v, u*) with high accuracy. Given this assumption, we expect that |*Ŵ* _*a,v*_ − *W*_*a,v*_| *>* |*Ŵ*_*n,v*_ − *W*_*n,v*_| if *v* is a neighbor of *a* and *a* is truly anomalous. This is because the predicted weight *Ŵ* _*a,v*_ for the edge (*a, v*) should deviate considerably from the actual weight *W*_*a,v*_ if *a* is anomalous, whereas the predicted weight *Ŵ* _*n,v*_ for the edge (*n, v*) should be close to the actual weight *W*_*n,v*_ if *n* is normal.

To implement this idea, we proposed a model that learns to predict the normal values of edge weights. We expect the model to fail to predict anomalous values. In other words, since most nodes and their edges are normal, it should be possible to train a model that learns to predict the values of normal interactions. A model that will learn to generalize well should successfully predict the normal values and fail to predict abnormal values, resulting in higher error for these cases.

#### 2.2.1 Construction of edge weight estimator

To create the predictive function *P*_*f*_ (*v, u*), we first trained a model that estimates edge weights. We used node embedding to generate features representing nodes *v* and *u* to train *P*_*f*_ (*v, u*). Node embedding is a vector representation of nodes in a graph that is designed to capture the relationships between nodes [14]. Unlike structure-based feature sets that may be specific to certain types of networks, node embedding is a more general solution that can produce near-state-of-the-art performance without requiring additional handcrafted features.

We treated the network as an unweighted graph to generate the embedding vectors.^1^ To select the node embedding method, we evaluated five types of embeddings: node2vec [15], RandNE [16], GLEE [17], NodeSketch [18], and DeepWalk [19] [20] based on the implementation by Rozemberczki et al. [20].

Using the extracted embeddings, we then trained a weight estimator using three regressors: Random Forest [21], XGBoost [22], and LightGBM [23]. The performance of the weight estimators was measured by *R*^2^ and Mean Squared Error (MSE). Also, it is important to note that during the training of the weight estimator, it is crucial to avoid considerable overfit.^2^ If the model overfits and “memorizes” the edge values like an infinitely large decision tree, it will not be able to identify nodes that deviate from the norm, as there will be no deviations in the predictions.

#### 2.2.2 Anomaly detection features

Anomaly detection features are used to identify nodes in the network that behave differently from the norm. Each feature represents a node by aggregating values representing the difference between the estimated and the actual edge weight of its edges. A larger average error is more likely to be anomalous, indicating that it behaves differently. The accumulation indicates that the deviation is not a result of the model error but something systematic. In the context of our study, we used anomaly detection features to identify proteins with important tissue-specific roles and interact differently with other proteins.

Deviation from the norm can be measured in a variety of ways. Similarly to Kagan et al. [13], we have generated anomaly detection features based on basic statistics such as STDV, mean, etc. To create the anomaly detection features, we defined the error of an edge (*u, v*) as *e*_*u,v*_ = *Ŵ* _*u,v*_ − *W*_*u,v*_ and the error of a node *v* as *e*_*v*_ = { *e*_*u,v*_ | *u* ∈ Γ(*v*) }. Based on these scores, we calculated 10 meta-features for each node in the network:

1. **Mean Error** - mean error of the predicted weights of the interactions between vertex *v* and its neighbors: 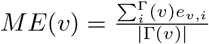.
2. **Error Standard Deviation** (Error SD) - dispersion of the predicted weights of the interactions between vertex *v* and its neighbors, *σ*(*e*_*v*_),
3. **Median Error** - median error of the predicted weights of the interactions between vertex *v* and its neighbors, *median*(*e*_*v*_).
4. **Sum of Errors** - the total sum of errors of the predicted weights of the interactions between vertex *v* and its neighbors, 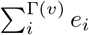.
5. **Standard error of the mean (SEM)** - measures the average amount of variation, 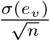.
6. **Mean Absolute Error** - absolute mean error of the predicted weights of the interactions between vertex *v* and its neighbors: 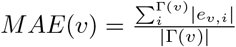.
7. **Absolute Error Standard Deviation** - dispersion of the absolute errors of the predicted weights of the interactions between vertex *v* and its neighbors, *σ*(| *e*_*v*_ |).
8. **Median Absolute Error** - the median of vertex *v* interaction prediction absolute errors: *median*(| *e*_*v*_ |).
9. **Sum of Absolute Errors** - the total sum of absolute errors for vertex *v*: ∑_*i*∈Γ(*v*)_ |*e*_*i*_ |.
10. **Standard Absolute Error of the mean** - measures the absolute average amount of variation: 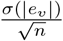.

To inspect the potential of the proposed features, we treated the extracted meta-features as a ten-dimension vector. Then, we used PCA to reduce the features into a two-dimensional vector. Next, we drew the PCA transformed data as a two-dimensional plot (see Fig. 2 and Supplementary information). In Fig. 2, we observed that most of the red points (anomalies) were grouped in the left area. This indicates that an anomaly detection algorithm can automatically identify these points.

**Figure 2:**
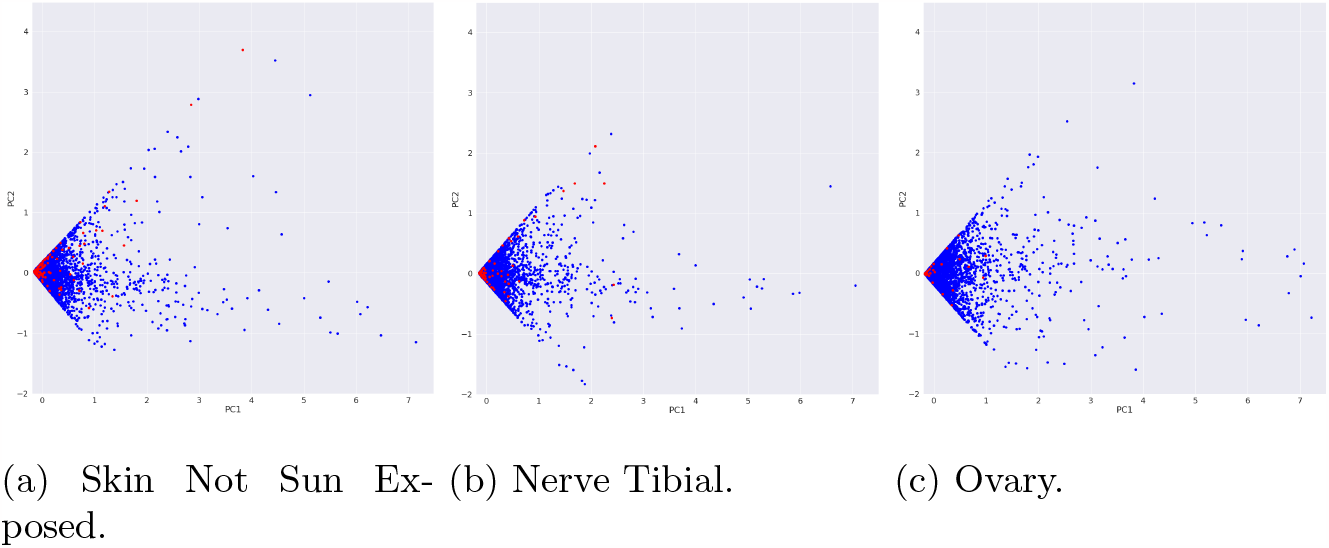
PCA representation of the anomaly detection features. The red dots represent disease-related proteins; the blue dots represent all other proteins.

#### 2.2.3 Anomalous nodes detection

The main approach to search for anomalies is by ranking the records according to a specific anomaly detection feature. Items with the highest values are considered potential anomalies. However, it is difficult to determine which of the 10 presented features is the most suitable for a specific use case. We propose four methods to deal with this issue to combine the generated anomaly detection features into one anomaly score.

1. **Feature’s Mean** - uses a sigmoid function to normalize the values of each feature, then assigns each node with the mean normalized values of the features. This produces a single value that can be used as the anomaly score.
2. **Principal Component Analysis (PCA)** - reduces the dimensionality of the ten features. The resulting component value can be used as the anomaly detection score [24].
3. **Isolation Forest (IForest)** - uses anomaly detection features as the IForest [25] model input to generate an anomaly score.
4. **Ensemble** - combines the previous three methods. Here, the raw Feature’s Mean, PCA values, and also their values after a sigmoid function were inserted into an IForest model to generate the anomaly score. Then, we select the max value from this new anomaly score and the scores produced from the first two methods.

### 2.3 Evaluation scheme

To assess the effectiveness of our method and evaluate anomaly detection performance, we performed a metrics-based evaluation. We measured the *Area under the ROC Curve* (*AUC*) [26] and *precision at k* (*P* @*K*), where *k* refers to the number of top-ranked records to consider. *AUC* is a widely used metric to estimate the overall performance of a classifier [26]. *Precision at k* is particularly useful in anomaly detection tasks, where we aim to discover new items and recommend or focus on the subset of predictions with the highest confidence scores. Based on these metrics, firstly, we compared the performance of five different node embedding models on weighted networks (see Section 2.2.1). Secondly, using the best-performing embedding model, we compared the performance of three edge weight estimator models, where their performance was also measured by *R*^2^ and mean squared error (MSE) (see Section 2.2.1). Thirdly, we used the best-performing combination of node embedding and edge weight estimator models and evaluated the performance of a variation of four different anomaly detection methods (see Section 2.2.3).

To evaluate the performance of the proposed method, we used two baselines:

1. **Node2Vec + IForest** - Similarly to Lee et al. [4], we used node2vec combined with IForest model. Here, the embeddings were generated based on the weights of the graph edges via node2vec. Then, these embeddings were used as the features for the IForest model.
2. **OddBall** - We used the OddBall algorithm [5] for anomaly detection in weighted graphs. To the best of our knowledge, OddBall is the only available algorithm with an implementation^3^ that was specifically designed for anomaly detection in weighted graphs.

### 2.4 Case Study

We illustrate the applicability of our method to the bioinformatic domain by using PPI networks with ground truth anomalies. In this use case, we expect that proteins involved in major biological roles in a specific tissue would have more anomalous interactions than other proteins in PPI networks of other tissues.

#### 2.4.1 Dataset of weighted tissue-specific PPI networks dataset

The dataset contains 17 PPIs signed networks (see Table 1) that were created by Basha et al. [27]. Each network was composed of 13,523 proteins (nodes) and 134,223 PPIs (edges). The weight of an edge ranged between [-1,1] and reflected whether the protein interaction was more likely (positive value) or less likely (negative value) to occur in that tissue relative to other tissues.

**Table 1:** PPI Networks Datasets.

| Tissue | #Disease-related nodes (%) |
| --- | --- |
| Artery Aorta | 23 (0.17) |
| Brain Cerebellum | 65 (0.48) |
| Brain Cortex | 77 (0.57) |
| Brain Spinal cord cervical c 1 | 48 (0.35) |
| Heart Atrial Appendage | 133 (0.98) |
| Heart Left Ventricle | 138 (1.02) |
| Liver | 53 (0.39) |
| Lung | 274 (2.03) |
| Muscle Skeletal | 121 (0.89) |
| Nerve Tibial | 77 (0.57) |
| Ovary | 41 (0.3) |
| Pituitary | 23 (0.17) |
| Skin Not Sun Exposed Suprapubic | 125 (0.92) |
| Skin Sun Exposed Lower leg | 125 (0.92) |
| Testis | 101 (0.75) |
| Whole Blood | 442 (3.3) |
| Whole Brain | 564 (4.2) |

#### 2.4.2 Annotation of disease-related proteins and tissue-specific biological processes

Data on Mendelian diseases, disease-related proteins, and disease-affected tissues were obtained from Hekselman et al. [28]. Nodes of a given tissue PPI network were labeled disease-related if the corresponding protein was associated with a Mendelian disease that affected that tissue. As shown in Table 1, up to 4.2% of the nodes per network were labeled disease-related. Data of proteins involved in additional tissue-selective diseases and phenotypes were obtained from PUBMED [29], GeneCards [30], and HPO [31] databases. Data of proteins participating in tissue-specific biological processes were obtained from Basha et al. [27]. Data of estimated activities of biological processes per protein and tissue were obtained from Sharon et al. [32]. Data of tissue risk assessment per protein, reflecting the likelihood of being disease-related in a given tissue, was available for 11 tissues from the TRACE webtool [33]. The 11 tissues were mapped to 13 tissue PPI networks (e.g., skin tissue was mapped to both sun-exposed and unexposed skin tissues).

### 2.5 Evaluation

#### 2.5.1 Edge weight prediction error

We tested whether predicted edge weights could be used to identify disease-related nodes. Per tissue, we constructed a tissue-specific PPI subnetwork where we only considered edges with high differences between the actual and the predicted values by RandNE estimator. Specifically, we only kept edges where the difference between the actual and predicted values was in the 95 quantiles. We hypothesized that nodes that appear on paths between disease-related nodes are highly likely to have central roles in the corresponding tissue. Hence, we counted how many times each node appeared on a path between two disease-related nodes, and considered the top-10 most frequent nodes per tissue PPI network. Next, we ranked nodes by their tissue risk assessment score, as computed by the TRACE webtool [33]. We compared the rank of the top-10 nodes of a tissue to their median rank in other tissues using a paired Wilcoxon test. In addition, per tissue, we compared the ranks of the top-10 nodes to the ranks of other nodes that were not included in the top-10 or were not disease-related using the Mann-Whitney test. P-values were adjusted for multiple hypothesis testing using the Benjamini-Hochberg procedure [34].

#### 2.5.2 Protein anomaly detection model

We tested whether the WGAND algorithm could be used to identify anomalous nodes with tissue-specific roles. We considered a protein to have a tissue-specific role if it was (i) designated as disease-related in that tissue or was involved in additional tissue-selective diseases and phenotypes, (ii) participated in a biological process that was specific to that tissue, or (iii) participated in a biological process that was more active in that tissue relative to other tissues. We applied WGAND to each tissue-specific PPI network, and focused on the top-10 anomalous nodes per network. To test for (i), we compared the fraction of disease-related proteins among the top-10 anomalous nodes relative to that fraction among all other nodes. To test for (ii), we computed the fraction of top-10 proteins that participated in a tissue-specific biological process out of the total number of proteins participating in such processes in the given tissue, and assessed for enrichment using the Fisher exact test. To test for (iii), we compared the estimated biological process activities of the top-10 anomalous proteins in the given tissue to their estimated activities in all other tissues using the Mann-Whitney test.

## 3 Results

To create an anomaly detection model based on weighted graphs, we used 17 tissue-specific PPI networks. We started by evaluating five different node embedding models and assessing their performance in detecting anomalous nodes via AUC, P@K, and runtime (see section 2.3). On average, RandNE embedding method showed the highest performance by all metrics, including the fastest average runtimes in generating the embeddings (see Table 2). Although its AUC is only slightly higher than other models, its P@K shows considerable gains against the rest of the models.

**Table 2:** Evaluation of different node embedding algorithms.

| Embedding | AUC | P@1 | P@3 | P@10 | P@20 | Embedding Runtime (Sec) |
| --- | --- | --- | --- | --- | --- | --- |
| DeepWalk | 0.6629 | 0.4118 | 0.2353 | 0.1941 | 0.1588 | 96 |
| GLEE | 0.6699 | 0.3529 | 0.2549 | 0.1765 | 0.1412 | 4 |
| Node2Vec | 0.6658 | 0.4118 | 0.2745 | 0.2412 | 0.1824 | 2912 |
| NodeSketch | 0.6700 | 0.4118 | 0.3137 | 0.2471 | 0.1941 | 229 |
| RandNE | **0.6701** | **0.5294** | **0.3725** | **0.2529** | **0.2147** | **1.6** |

Next, we tested three different edge weight estimators and evaluated their performance (see section 2.3). First, we evaluated their performance just on the edge weight estimation task. In terms of MSE LightGBM had the best performance (see Table 3a). Afterward, we evaluated the performance of the whole anomaly detector using the different weight estimator models as backbones (see Table 3b). We found that on average when using the RandomForest edge weight estimator to generate the features, the anomaly detection model achieved the highest score in all anomaly detection metrics. The rest of the results were based on the combination of RandNE embedding and RandomForest edge weight estimator, which presented the best results. We turned to check whether the edge weight estimator could provide biological (see Section 2.5.1). We created a PPI subnetwork that only contained edges with a high difference between the predicted value and the actual value per tissue network. We focused on the top-10 proteins (nodes) per network that participated in the largest number of paths between two disease-related proteins. We used the TRACE tool [35] to rank of the proteins by on their likelihood of being disease-related in the given tissue. The top-10 proteins were ranked significantly higher in the given tissue relative to their median ranking in other tissues (*p* − *value* = 7.57*e* − 6, Wilcoxon test; Fig. 3A). In addition, except for brain cerebellum tissue, the top-10 proteins ranked significantly higher relative to other proteins of the same tissue (adjusted *p* − *value* ≤ 8.12*e* − 3, Mann-Whitney test, Benjamini-Hochberg correction; Fig. 3B). These results show that the top-10 proteins were not randomly picked but could potentially have important tissue-specific roles. Overall, this analysis suggests that based on the error in predicting the edge weight, we can identify anomalous nodes.

**Table 3:**
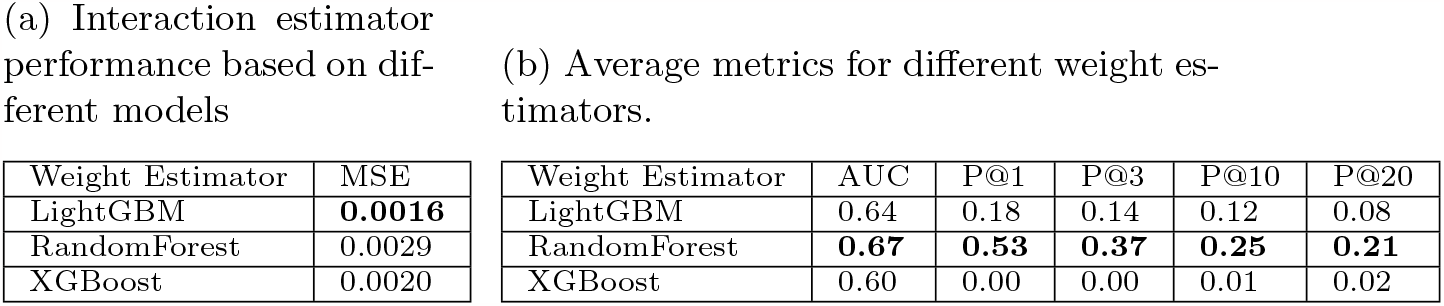
Evaluation of different edge weight estimation models.

**Figure 3:**
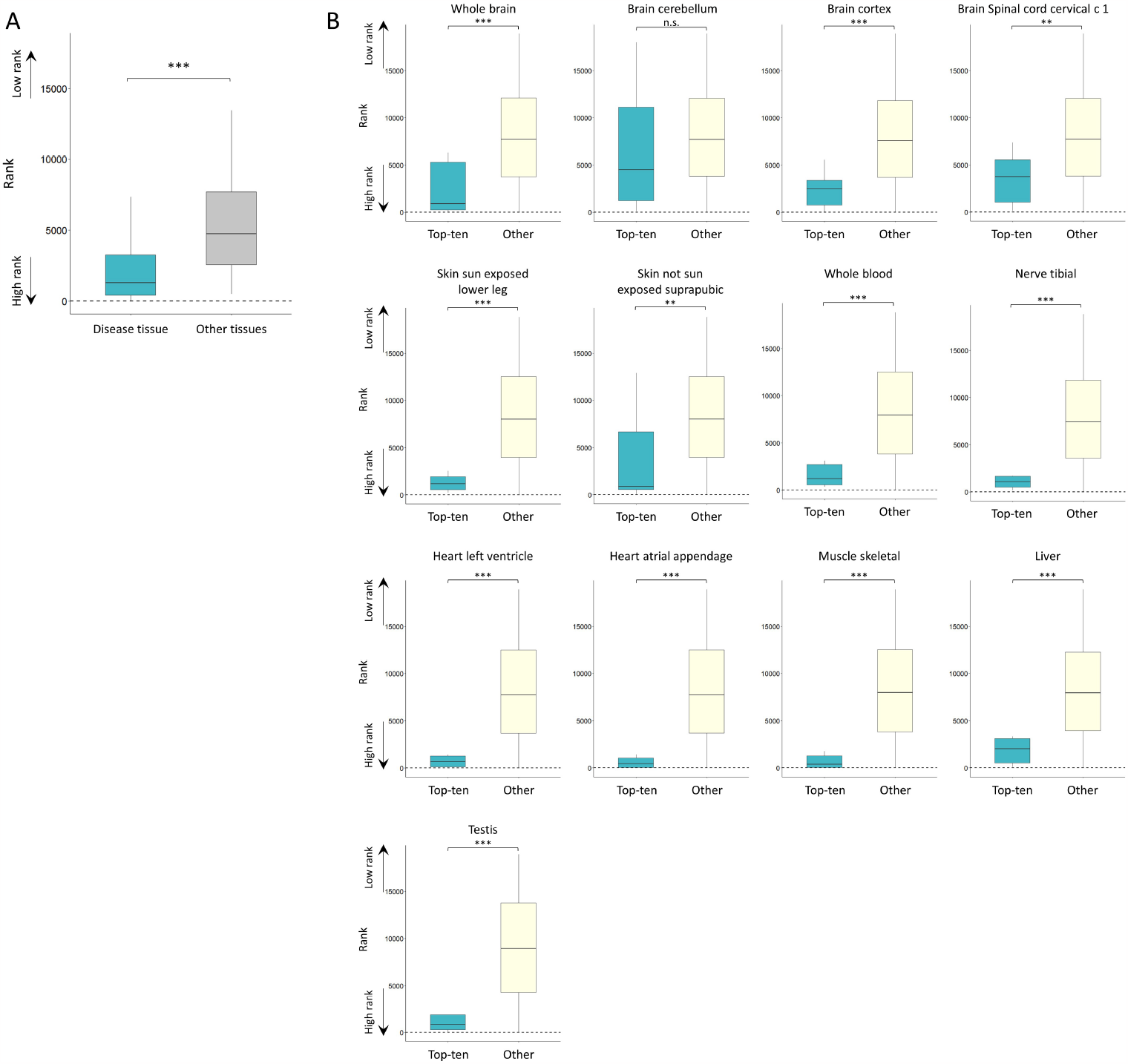
Pathway analysis validation. **A)** Top-10 proteins of a given tissue network were significantly more likely to be disease-related in that tissue (blue) than in other tissues (grey; p-value=7.57e-6, Wilcoxon test). Top-10 proteins were collated from 13 tissue networks. **B)** Top-10 proteins (blue) were significantly more likely to be disease-related in the given tissue than other proteins (beige). Adjusted p-values for whole brain, cerebellum, cortex, spinal cord, skin sun-exposed, skin not sun-exposed, whole blood, nerve, heart left ventricle, heart atrial appendage, muscle, liver, and testis in respective order: 8.6e-4, 0.13, 8.14e-4, 0.081,9.83e-5, 1.85e-3, 1.16e-4, 3.85e-5, 2.46e-6, 2.75e-6, 3.03e-5, 9.21e-4 and 3.11e-5 (one-sided Mann-Whitney test, Benjamini-Hochberg correction).

Next, we aimed to construct an unsupervised machine-learning-based anomaly detector to detect anomalous nodes. Per tissue PPI network, we assigned network nodes with ten meta-features that were constructed based on the edge weight prediction error (see Section 2.2.2). Then, using the ground truth labels, we evaluated the performance of four suggested anomaly detector models and compared them to two baseline models (see Section 2.3). On average, all four anomaly detector models present superior performance over the baseline models in all metrics with respect to AUC and P@K (see Fig. 4). In terms of AUC, the ensemble and the feature’s mean presented the best performance with a marginal difference. In terms of P@K, we saw similar performance between the ensemble, PCA, and the feature’s mean with a slight advantage for the ensemble. In addition, we evaluated the performance of the ensemble method against the baseline models of each tissue network separately. The ensemble method had a higher AUC than the baselines across all tissues and higher P@K in 13 out of 17 of the tissues (see Table S1). Overall, the ensemble method – ‘WGAND (ensemble)’ - showed the best performance. For 85% of the datasets, the proposed method achieved higher performance than the baselines in terms of AUC and P@K. We noticed in Figures 5 and S1 that there is a significant variance between the different methods in most tissues. Also, we noticed that tissue where the anomaly detector achieves high AUC but does not always translate into high P@K.

**Figure 4:**
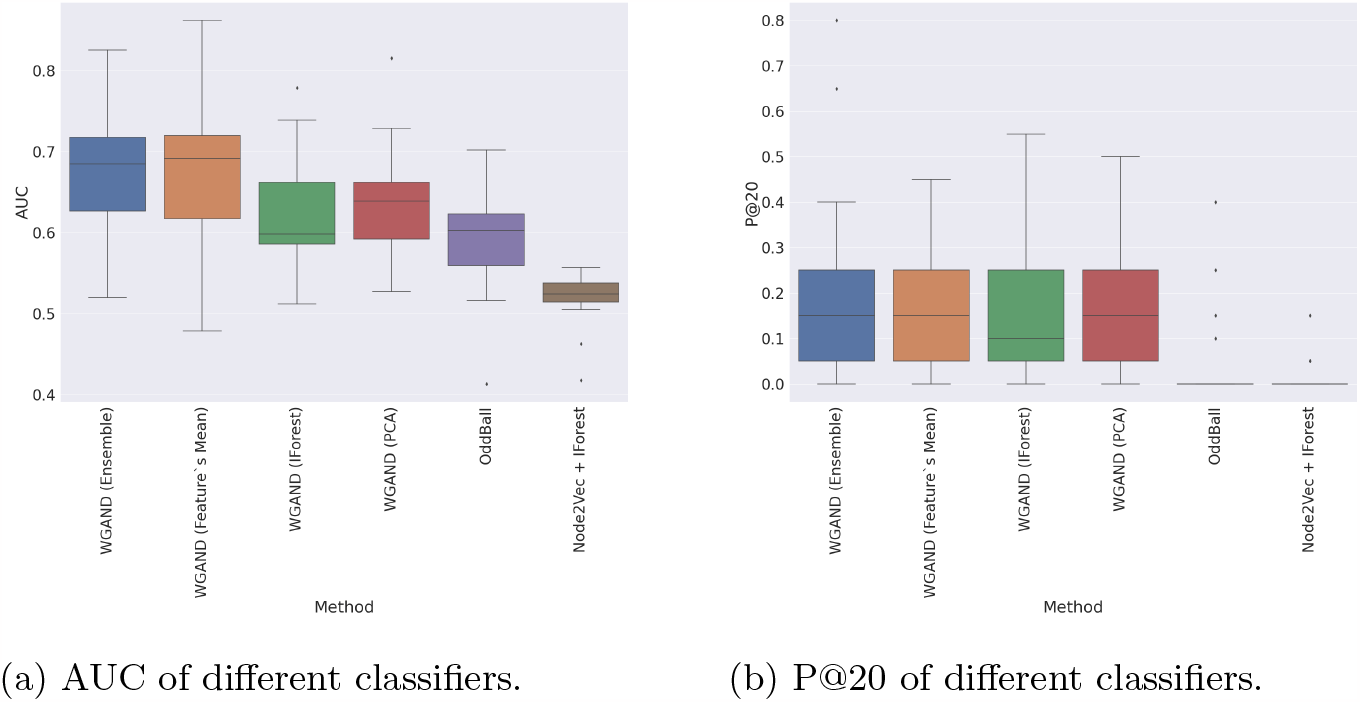
Evaluation of different classifiers to predict anomalous nodes based on the meta-features.

**Figure 5:**
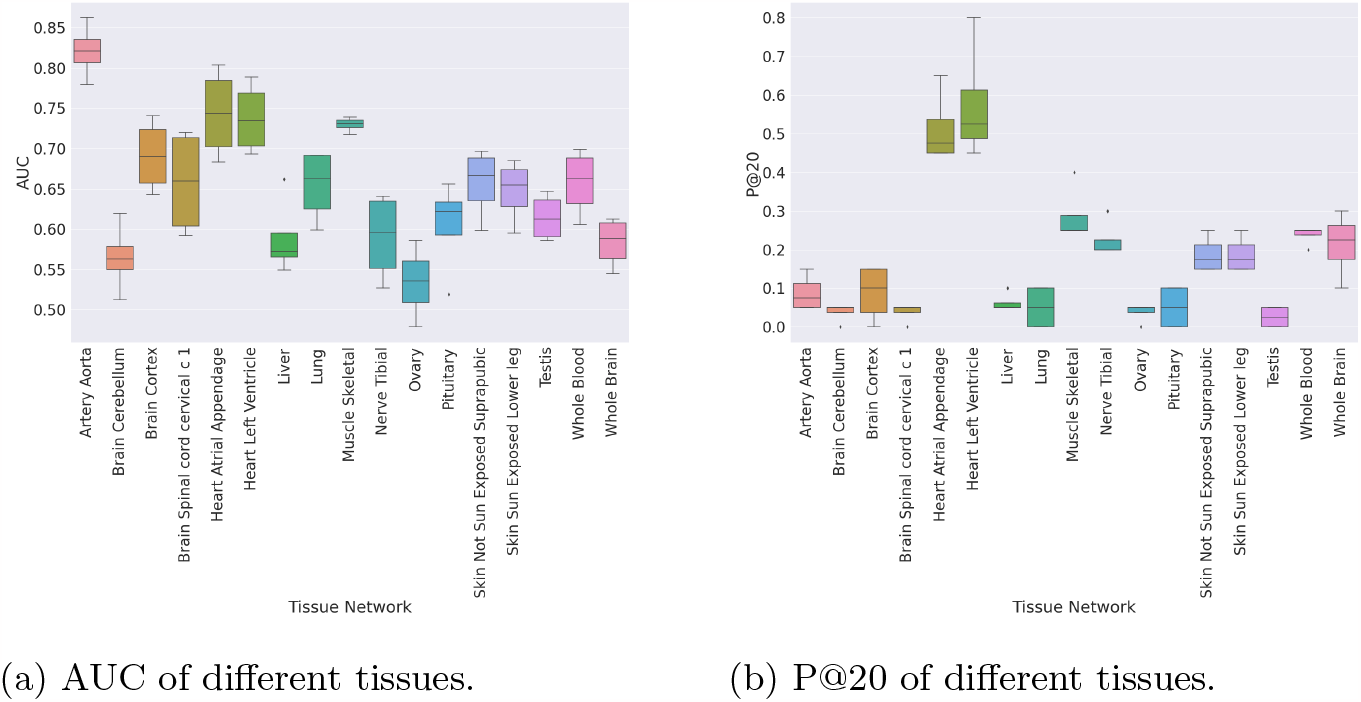
Evaluation of the variance of the four methods for predicting anomalous nodes for each tissue. We can observe that from most tissues, there is a high variance in performance between different methods.

Next, to inspect which feature could explain anomaly best and may be used as an indicator in unlabeled networks, we used the WGAND ‘ensemble’ method and constructed a model per feature. The models were evaluated based on their P@K values (see Figs. S2). Most features had P@K higher than random in all the tissue networks (see Table S2). The features with the best performance on average were the *Sum of Errors, Error SD*, and *Mean Error*.

We hypothesized that tissue-associated anomalous proteins would play major biological roles in that tissue, which would be evident either by their involvement in a disease that affected that tissue or in tissue-specific or tissue-preferentially-active biological processes (see Section 2.5.2). We tested this hypothesis for the top-10 anomalous proteins per tissue network according to the best-performing combination “WGAND (ensemble).” First, we focused on the involvement of these proteins in Mendelian diseases. We found that 26% of the top-10 anomalous proteins per tissue (a total of 170 proteins) annotated as disease-related in that tissue [28], relative to 1.5% when considering all proteins per tissue (Fig. 6A). An additional 24% of the top-10 proteins were associated with tissue-selective diseases and phenotypes according to literature and disease-related databases (Fig. 6 and Table S3). Hence, top-10 anomalous proteins were indeed highly enriched for proteins involved in diseases and phenotypes affecting that tissue. Next, we checked whether the top-10 anomalous proteins were involved in tissue-specific or preferentially active biological processes in the corresponding tissue. For that, we first calculated the overlap between the top-10 anomalous proteins and proteins involved in tissue-specific biological processes. The overlap was significant (fig. 5C; p-value=6.5e-49, Fisher Exact test). Moreover, the top-10 anomalous proteins were also more likely involved in processes that were active preferentially in the corresponding tissue (Fig 5C; p = 2.5e-16, Mann-Whitney test). We applied similar tests to the top-10 anomalous proteins that were detected by the other three WGAND (feature’s mean, IForest, and PCA). They also showed significant yet slightly weaker enrichments (Fig. 6).

**Figure 6:**
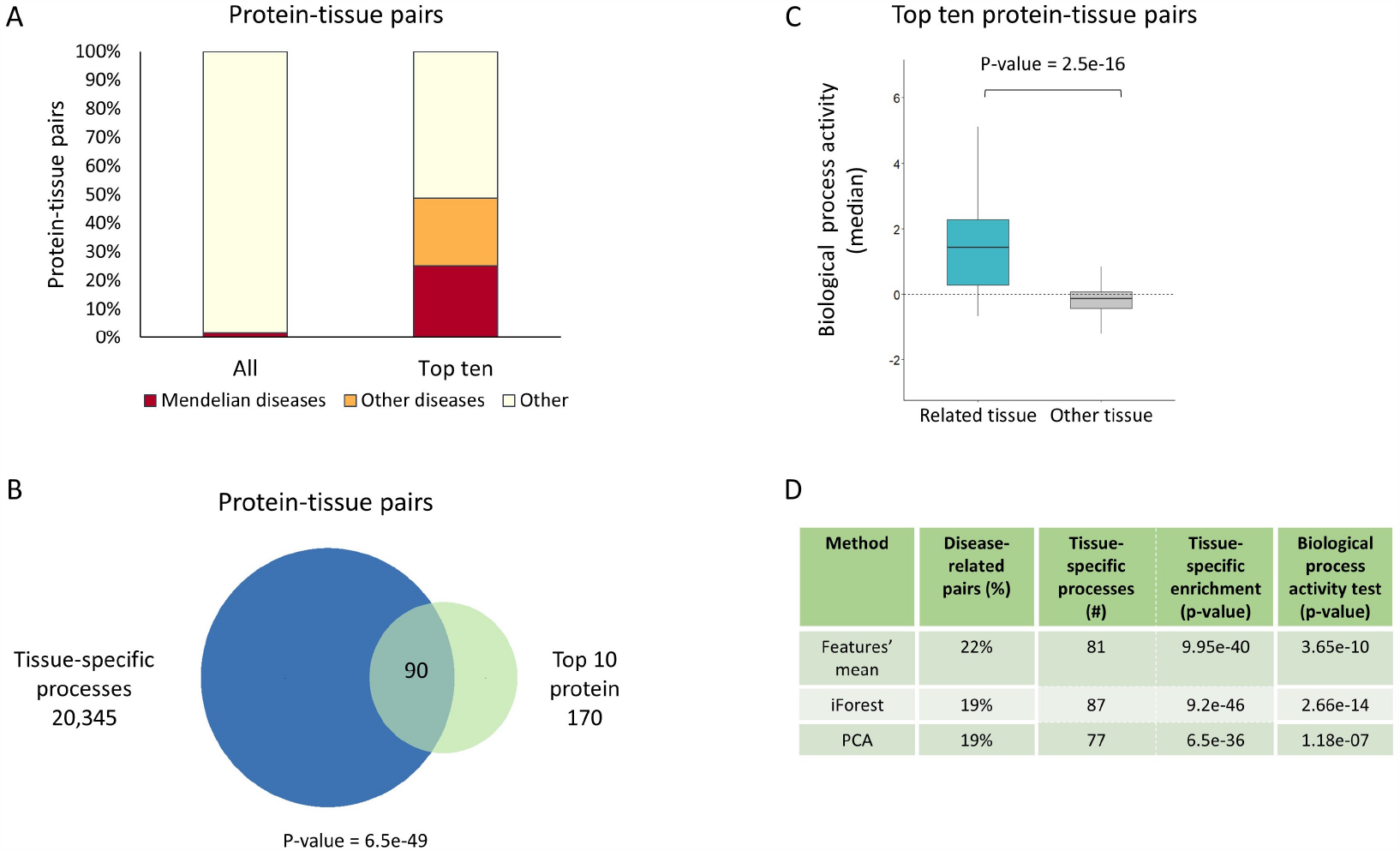
Top-10 anomalous proteins were enriched for tissue-specific diseases and cellular processes. (a) Percentage of protein-tissue pairs where the protein was associated with a disease affecting the corresponding tissue. Top-10 pairs were enriched for Mendelian disease-related proteins (26%, red) in contrast to all pairs (1.5%, red) and were also enriched for proteins involved in other tissue-selective diseases and phenotypes (24%, orange). (b) The overlap between the top-10 protein-tissue pairs (170 pairs) and pairs where the protein was associated with a biological process that was specific to that tissue (20,345 pairs). Top-10 pairs were enriched for tissue-specific processes (90 pairs, p-value = 6.5e-49, Fisher exact test). (c) Top-10 proteins per tissue were more likely associated with cellular processes that were preferentially active in that tissue (blue) relative to other tissues (grey; p-value = 2.5e-16, Mann-Whitney). (d) Analyses in panels (a-c) were applied to the top-10 anomalous proteins that were detected by three the methods including features’ mean, IForest and PCA. The results were significant yet weaker than the ensemble method.

In conclusion, our findings indicate that WGAND outperforms the baselines in detecting anomalous nodes. We have also shown, through PCA visualization, that the generated features were capable of distinguishing between anomalous and normal nodes. Furthermore, we demonstrated that the top ten anomalous proteins overlapped significantly with proteins involved in tissue-specific or tissue-preferentially-active biological processes.

## 4 Discussion

We presented a novel machine learning-based method for detecting anomalous nodes in signed and weighted networks, which has important implications for various domains. Here, we demonstrated the effectiveness of our approach by applying it to PPI networks of different human tissues to identify anomalous proteins that could have important biological roles in the respective tissues.

First, when dealing with imbalanced data, P@K is critical, as a model with higher P@K can help experts narrow down the search space when inspecting anomalies. While the differences between various node embedding models are not highly significant, it is possible that the performance differences could be more pronounced in different domains. We plan to investigate this further in future studies by analyzing additional node embedding models and networks.

We observed that a lower MSE for the edge weight estimator does not necessarily mean that the anomaly detection model would have a higher AUC or P@K. In fact, the RandomForest-based edge weight estimator had the highest MSE, yet it had the highest scores on average with all node classifiers. This could be due to overfitting by the other models, resulting in lower error. Additionally, the RandomForest might make mistakes on problematic interaction values, which could reduce its performance as measured by the metrics and make the error more evident when aggregated.

Also, we found that the WGAND ensemble performs significantly better than the baseline models in most tissues, with Oddball outperforming WGAND in only four brain-related networks in terms of P@K but worse in terms of AUC. This suggests that biological differences between brain tissues and other tissues may be a contributing factor to the observed performance differences. In future studies one could explore further the commonalities and differences between brain related tissues. Node2Vec + IForest was not able to achieve good results with this data in all the tissues suggesting that it was not suited for the task.

Next, we showed that a single meta-feature can be a good measure for node anomaly detection. On average, the Sum of Errors is the best predictive single feature across all metrics. Surprisingly, the Absolute Sum of Errors is the worst feature across all metrics, despite our expectation that the error direction should not matter and that a higher total error should be a good indication of an anomaly. This suggests that error direction is an important factor in node anomaly detection and that the difference between regular and absolute meta-features is inconsistent. In a future study, we would like to evaluate the WGAND on different use cases such as both detection and malicious users detection. We would like to explore if the same type of anomaly detection features would continue to give the best performance also in different domains.

Additionally, we investigated different methods for generating anomaly scores and extracting anomalies from the generated anomaly features and found that some methods have similar performance. Among the four proposed methods, the IForest-based solution performs the worst in terms of AUC, while Feature’s Mean has the best AUC. In terms of P@K, the differences are smaller, but different methods have advantages in different tissue networks. For example, the ensemble-based method has a significant advantage in the liver network, while the PCA-based and Feature’s Mean methods have a significant advantage in the lung network. Also, some tissues, such as Skin Not Sun Exposed Suprapubic, Heart Left Ventricle, and Nerve Tibial single features, had higher P@K than the machine learning model. While it is difficult to conclude which method is the best, a good rule of thumb would be first to try using the ensemble-based WGAND, which should combine the advantages of all other methods.

## 5 Data availability

The code and datasets generated during and analyzed during the current study are available in the project repository^4^.

## 6 Acknowledgements

This study was funded by the Israeli Council for Higher Education (CHE) via the Data Science Research Center, Ben-Gurion University of the Negev, Israel [to M.F. and E.Y.-L], and by the Israel Science Foundation [401/22 to E.Y.-L.]. J.J. wishes to thank the Baroness Ariane de Rothchild Women Doctoral Program.

## 7 Supplementary

**Figure S1:**
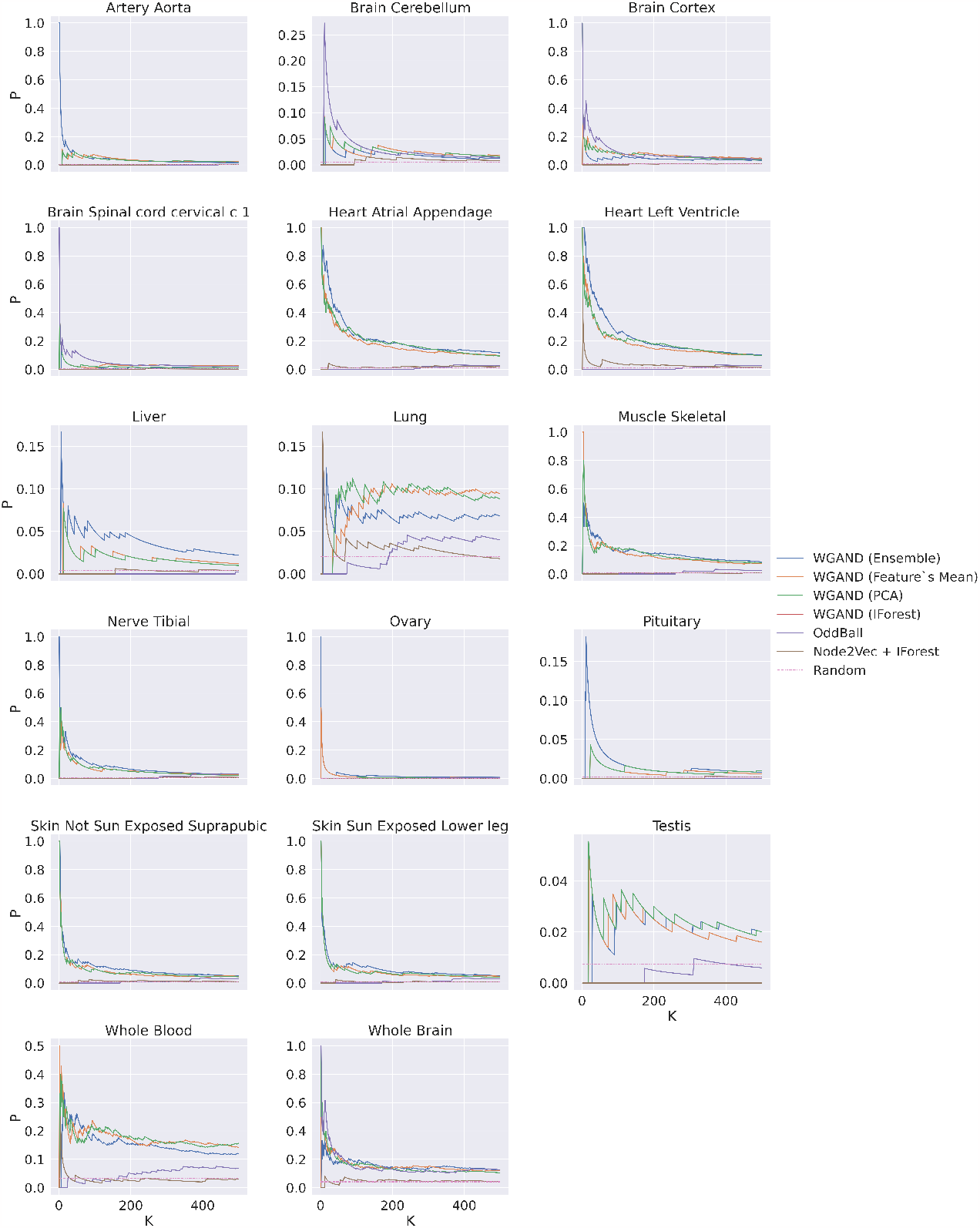
P@K of different classifiers per each PPI tissue-specific network.

**Figure S2:**
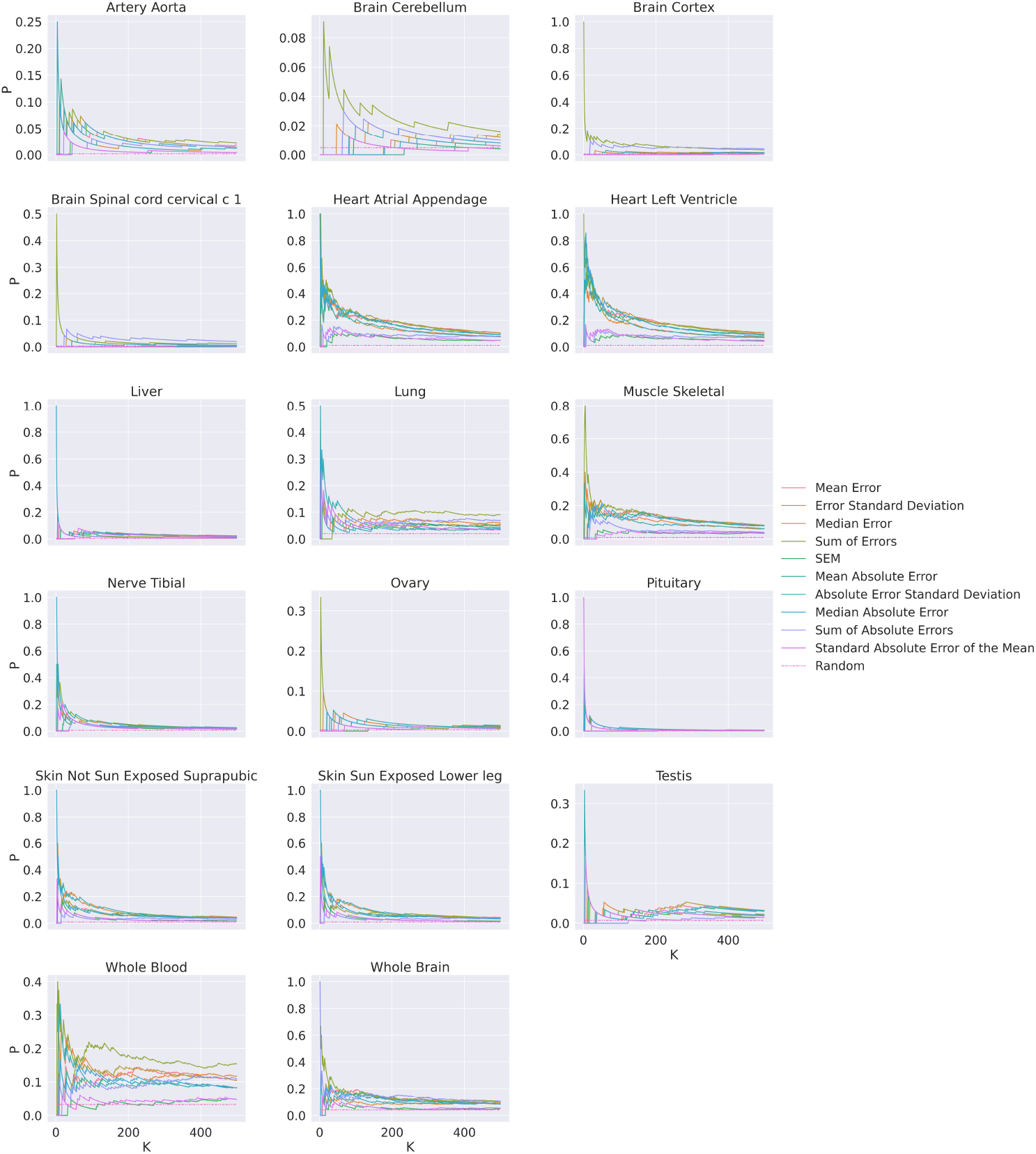
P@K of WGAND ‘ensemble’ method trained with each feature separately on PPI tissue-specific networks.

**Figure S3:**
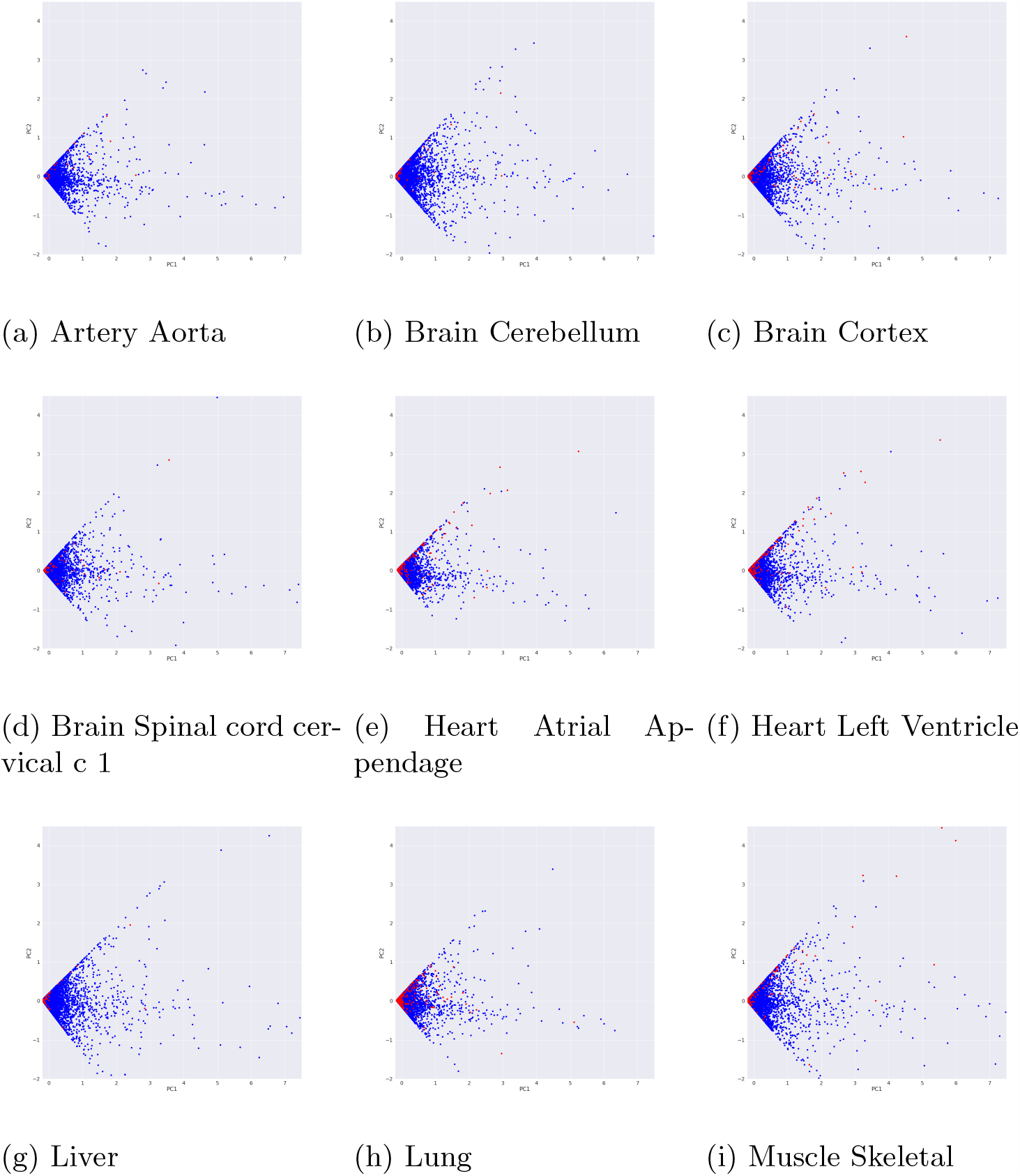
PCA representation of the anomaly detection features. The red dots represent disease-related proteins; the blue dots represent all other proteins.

**Figure S4:**
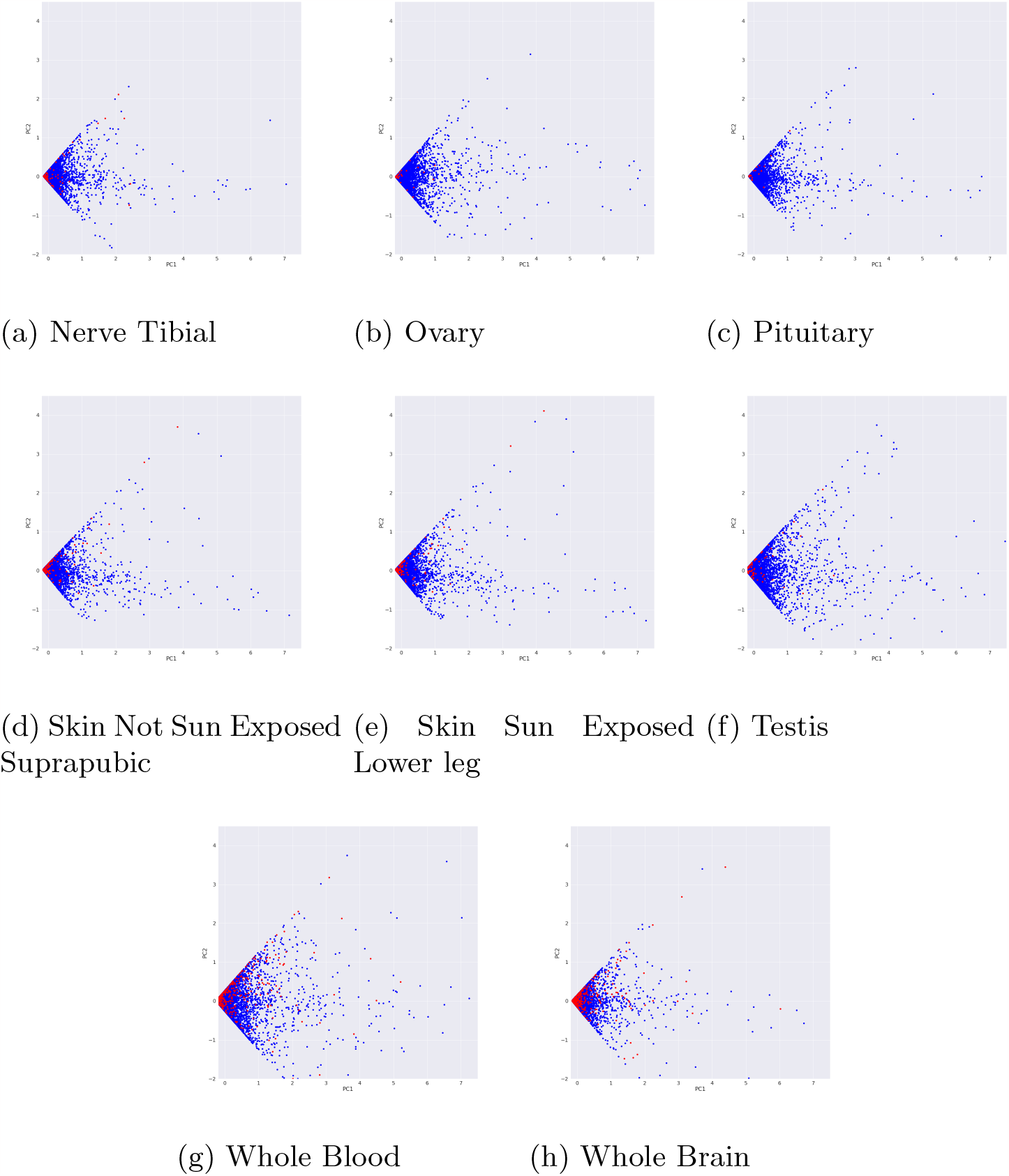
PCA representation of the anomaly detection features. The red dots represent disease-related proteins; the blue dots represent all other proteins.

**Table S1:** Evaluation WGAND versus baselines on PPI dataset.

| Tissue Network | WGAND (Ensemble) |  |  |  |  | Node2Vec + Iforest |  |  |  |  | Oddball |  |  |  |  |
| --- | --- | --- | --- | --- | --- | --- | --- | --- | --- | --- | --- | --- | --- | --- | --- |
|  | AUC | P@1 | P@3 | P@10 | P@20 | AUC | P@1 | P@3 | P@10 | P@20 | AUC | P@1 | P@3 | P@10 | P@20 |
| Artery Aorta | **0.83** | **1.00** | **0.67** | **0.20** | **0.15** | 0.46 | 0.00 | 0.00 | 0.00 | 0.00 | 0.70 | 0.00 | 0.00 | 0.00 | 0.00 |
| Brain Cerebellum | **0.56** | 0.00 | 0.00 | 0.00 | 0.05 | 0.52 | 0.00 | 0.00 | 0.00 | 0.00 | 0.53 | 0.00 | 0.00 | **0.20** | **0.15** |
| Brain Cortex | **0.72** | 0.00 | **0.33** | 0.10 | 0.05 | 0.53 | 0.00 | 0.00 | 0.00 | 0.00 | 0.68 | **1.00** | **0.33** | **0.40** | **0.25** |
| Brain Spinal cord cervical c 1 | **0.71** | 0.00 | 0.00 | 0.10 | 0.05 | 0.52 | 0.00 | 0.00 | 0.00 | 0.05 | 0.70 | **1.00** | **0.33** | **0.20** | **0.10** |
| Heart Atrial Appendage | **0.78** | **1.00** | **1.00** | **0.70** | **0.65** | 0.56 | 0.00 | 0.00 | 0.00 | 0.00 | 0.63 | 0.00 | 0.00 | 0.00 | 0.00 |
| Heart Left Ventricle | **0.76** | **1.00** | **1.00** | **0.90** | **0.80** | 0.54 | 0.00 | 0.00 | 0.00 | 0.00 | 0.62 | 0.00 | 0.00 | 0.00 | 0.00 |
| Liver | **0.55** | 0.00 | 0.00 | 0.00 | **0.10** | 0.42 | 0.00 | 0.00 | 0.00 | 0.00 | 0.41 | 0.00 | 0.00 | 0.00 | 0.00 |
| Lung | **0.69** | **0.00** | **0.00** | **0.10** | **0.10** | 0.52 | 0.00 | 0.00 | 0.10 | 0.05 | 0.60 | 0.00 | 0.00 | 0.00 | 0.00 |
| Muscle Skeletal | **0.73** | **1.00** | **0.67** | **0.50** | **0.40** | 0.54 | 0.00 | 0.00 | 0.00 | 0.00 | 0.56 | 0.00 | 0.00 | 0.00 | 0.00 |
| Nerve Tibial | **0.63** | **1.00** | **0.33** | **0.30** | **0.30** | 0.51 | 0.00 | 0.00 | 0.00 | 0.00 | 0.59 | 0.00 | 0.00 | 0.00 | 0.00 |
| Ovary | **0.52** | **1.00** | **0.33** | **0.10** | **0.05** | 0.51 | 0.00 | 0.00 | 0.00 | 0.00 | 0.52 | 0.00 | 0.00 | 0.00 | 0.00 |
| Pituitary | **0.63** | **0.00** | **0.00** | **0.20** | **0.10** | 0.52 | 0.00 | 0.00 | 0.00 | 0.00 | 0.53 | 0.00 | 0.00 | 0.00 | 0.00 |
| Skin Not Sun Exposed Suprapubic | **0.69** | **1.00** | **0.67** | **0.30** | **0.20** | 0.54 | 0.00 | 0.00 | 0.00 | 0.00 | 0.62 | 0.00 | 0.00 | 0.00 | 0.00 |
| Skin Sun Exposed Lower leg | **0.67** | **1.00** | **0.67** | **0.30** | **0.20** | 0.52 | 0.00 | 0.00 | 0.00 | 0.00 | 0.62 | 0.00 | 0.00 | 0.00 | 0.00 |
| Testis | **0.63** | **0.00** | **0.00** | **0.00** | **0.00** | 0.51 | 0.00 | 0.00 | 0.00 | 0.00 | 0.57 | 0.00 | 0.00 | 0.00 | 0.00 |
| Whole Blood | **0.69** | **1.00** | **0.33** | **0.20** | **0.25** | 0.54 | 0.00 | 0.00 | 0.10 | 0.15 | 0.61 | 0.00 | 0.00 | 0.00 | 0.00 |
| Whole Brain | **0.61** | 0.00 | 0.33 | 0.30 | 0.20 | 0.53 | 0.00 | 0.00 | 0.00 | 0.00 | 0.56 | **1.00** | **0.67** | **0.50** | **0.40** |
| Average | **0.67** | **0.53** | **0.37** | **0.25** | **0.21** | 0.52 | 0.00 | 0.00 | 0.01 | 0.01 | 0.59 | 0.18 | 0.08 | 0.08 | 0.05 |

**Table S2:**
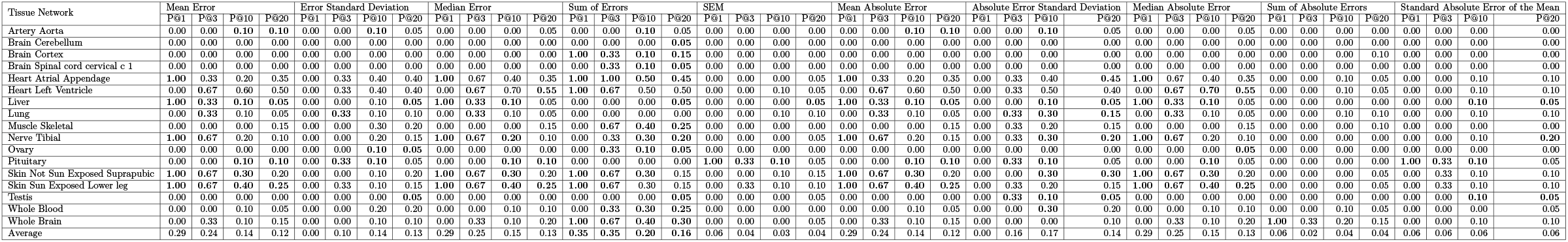
P@K of WGAND ‘ensemble’ method constructed per feature.

**Table S3:**
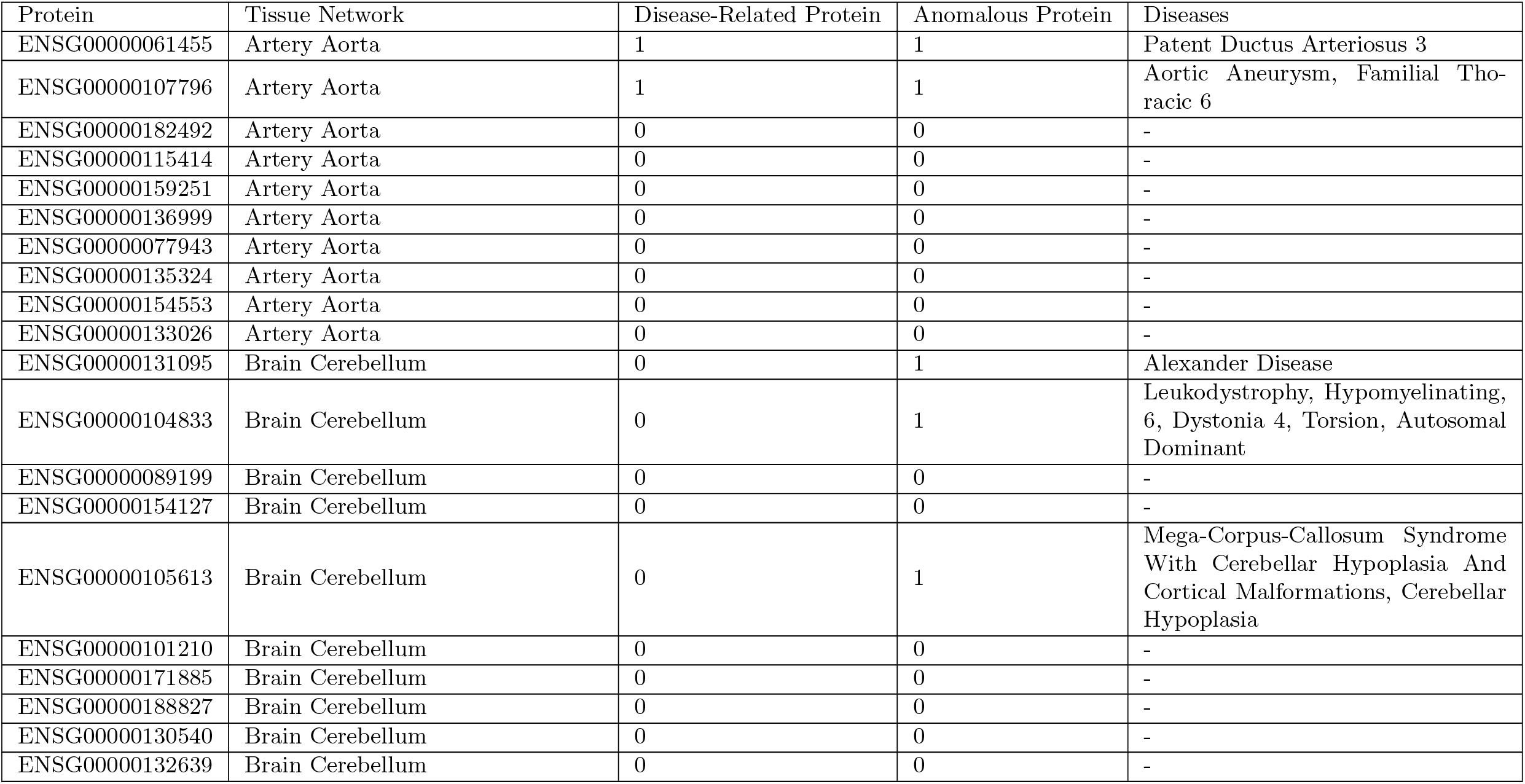
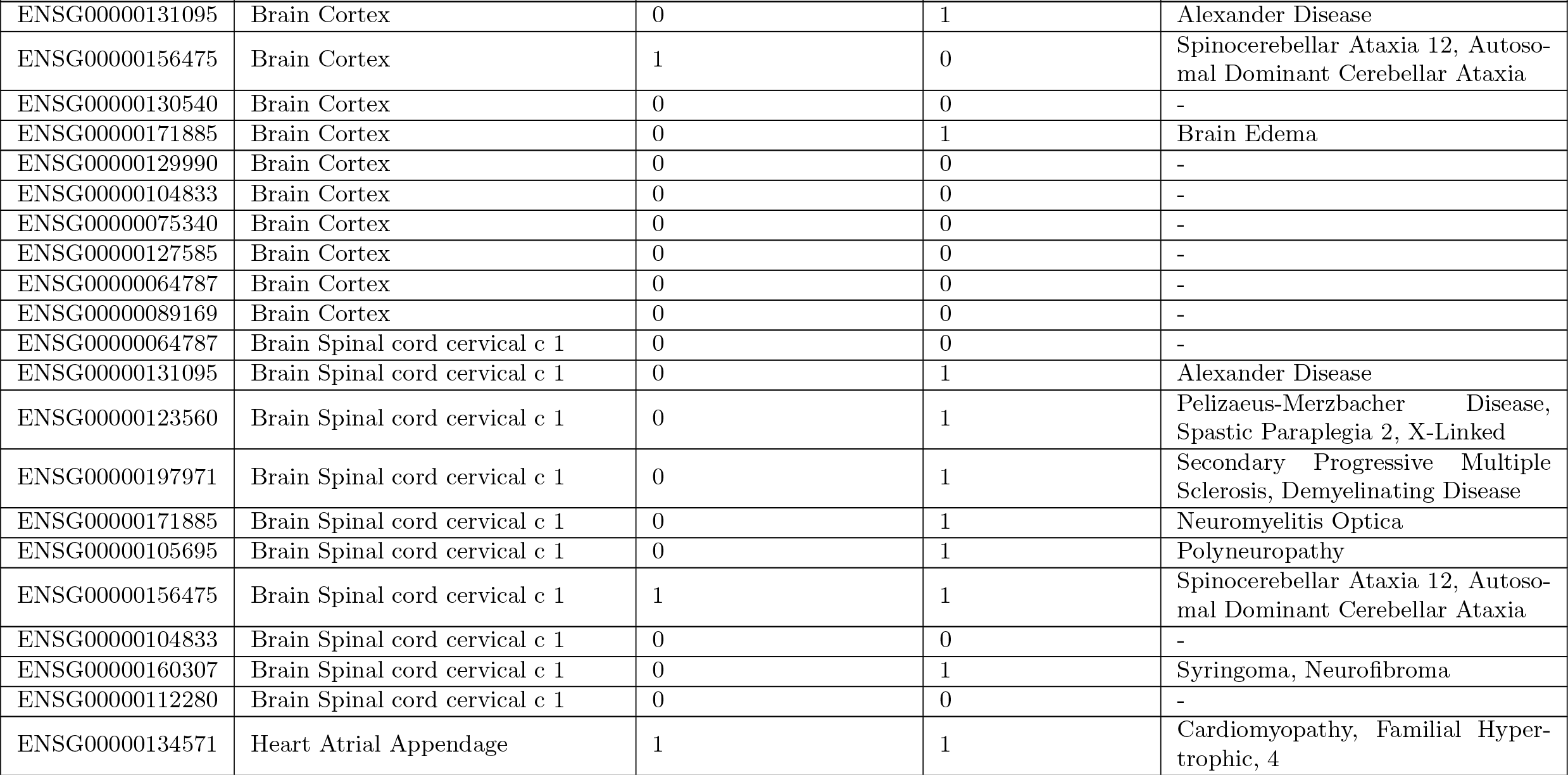
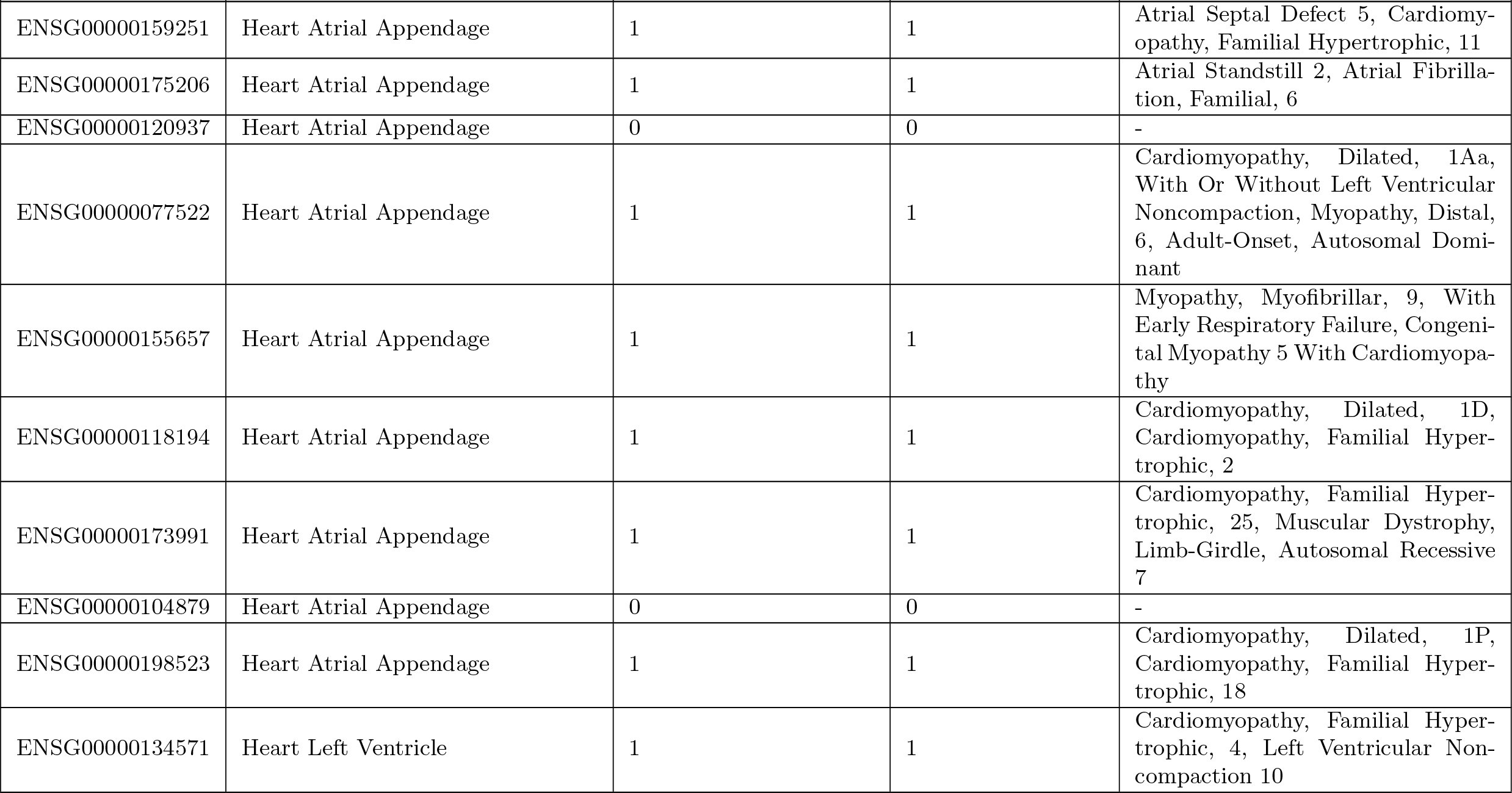
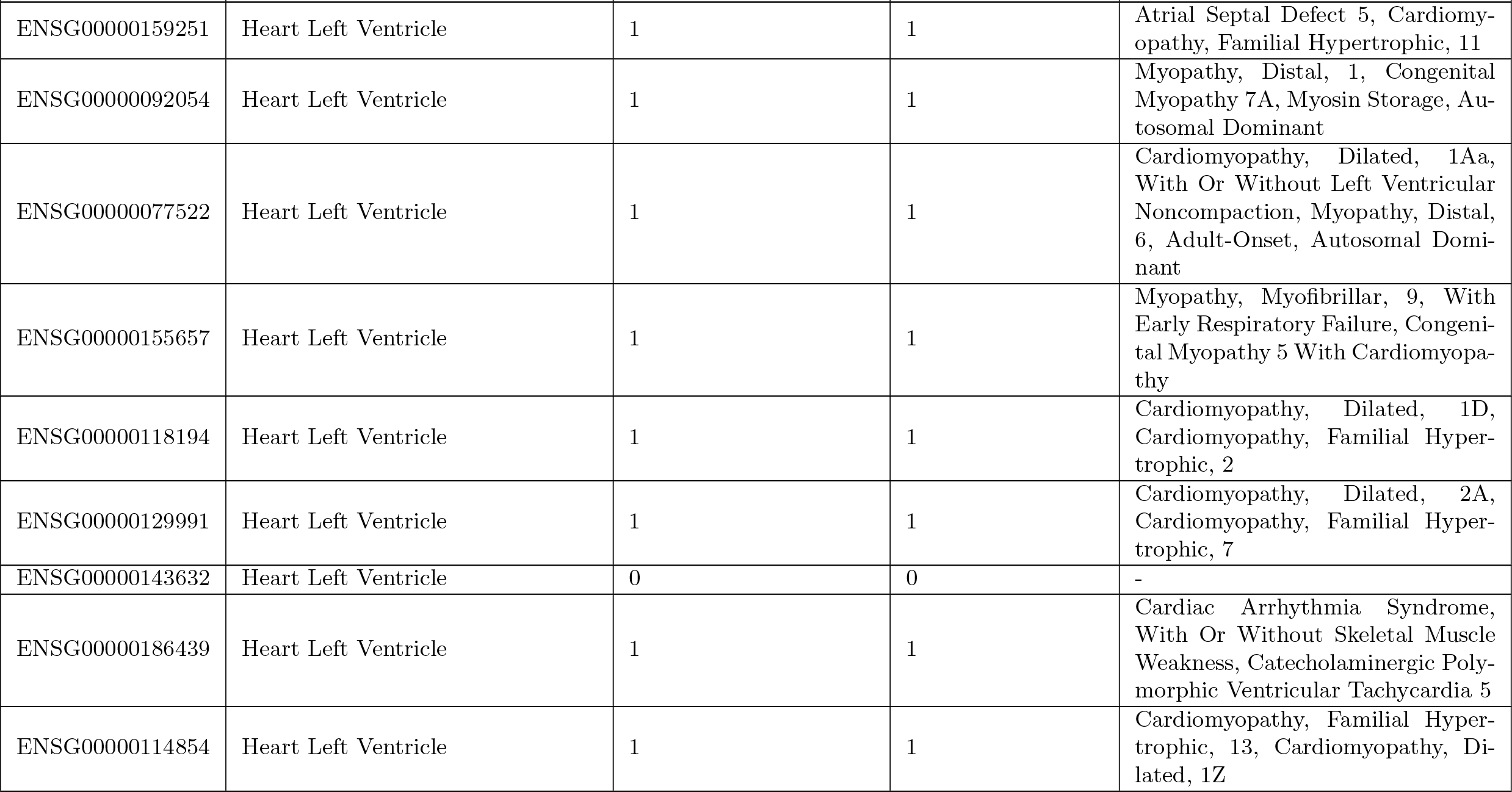
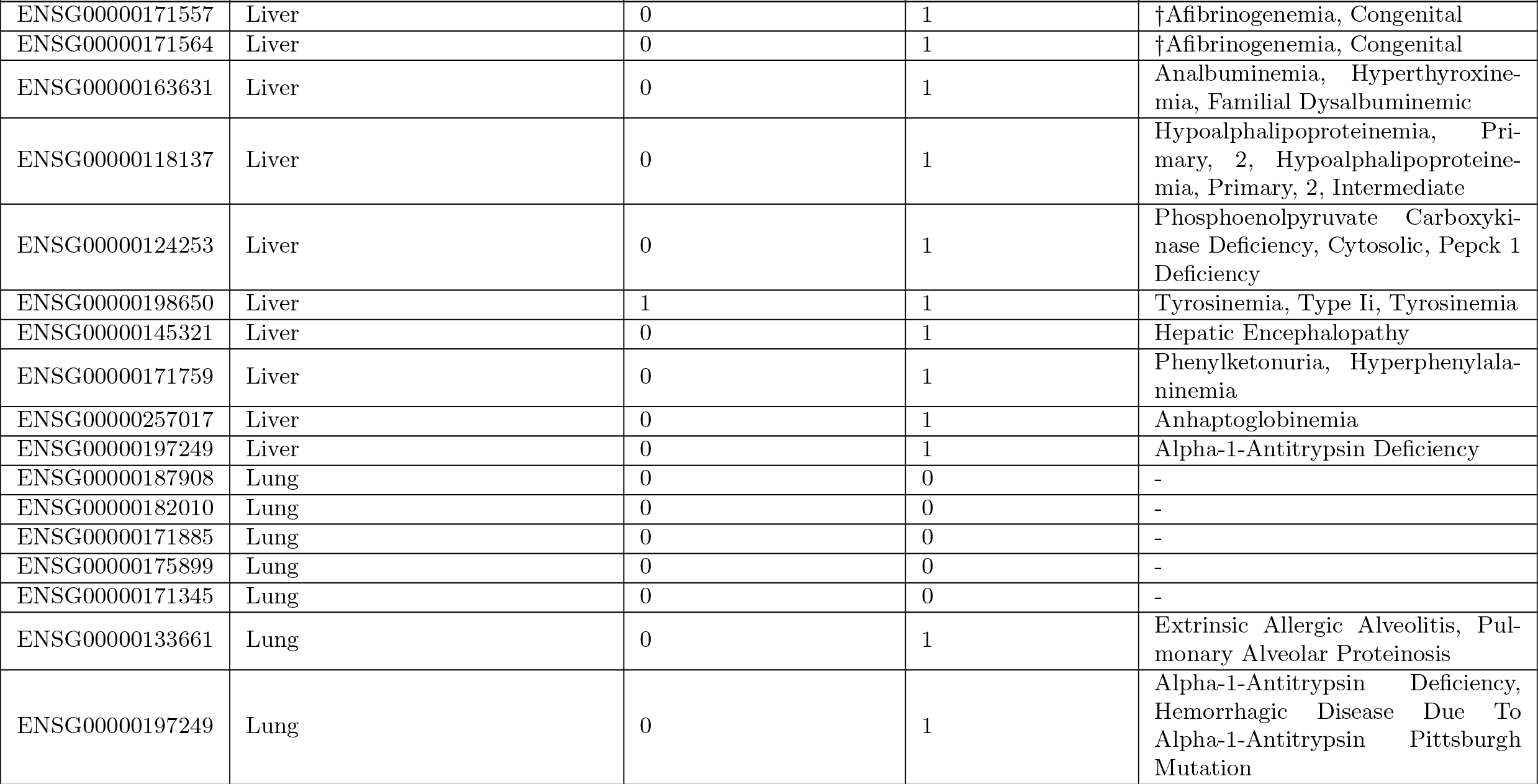
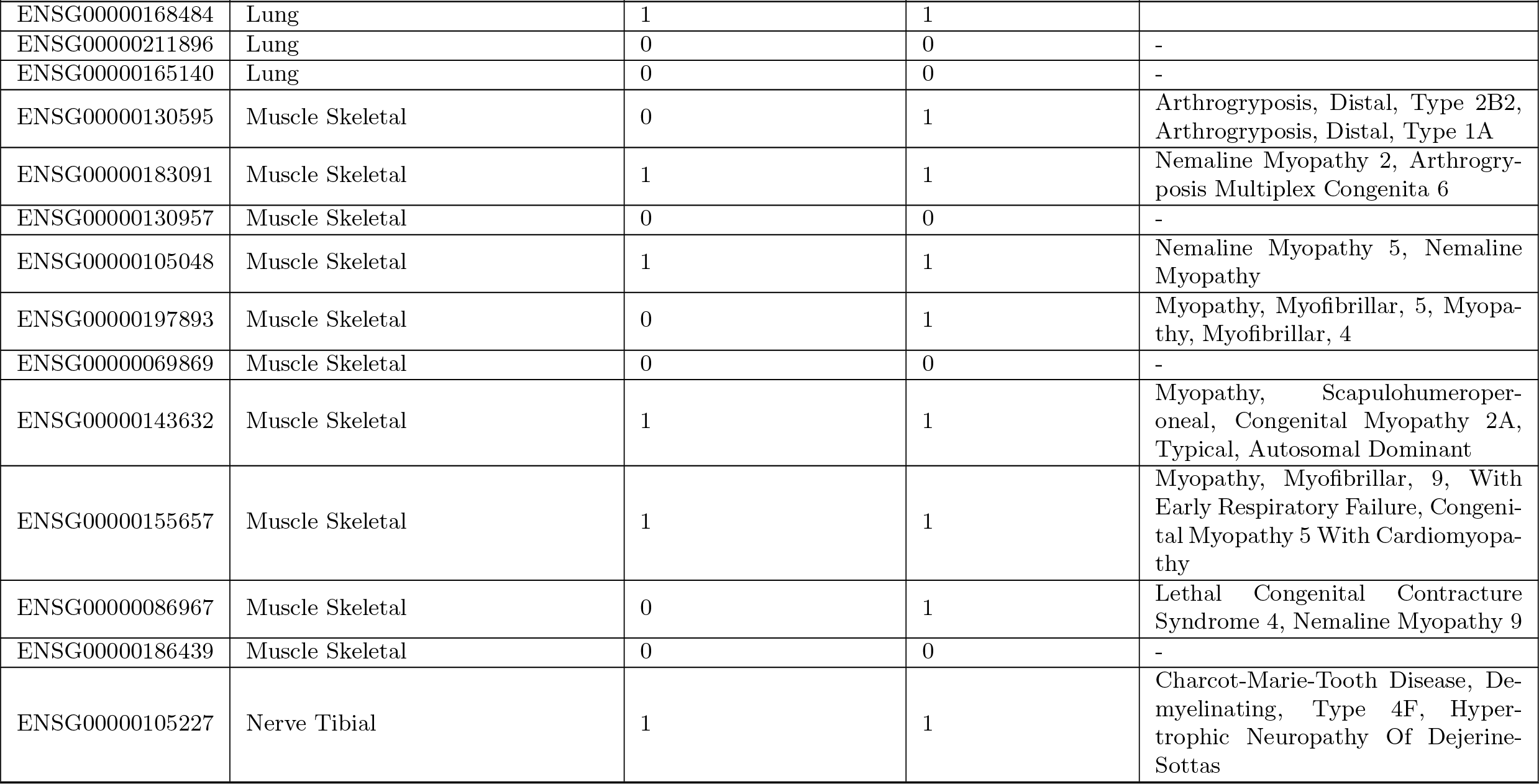
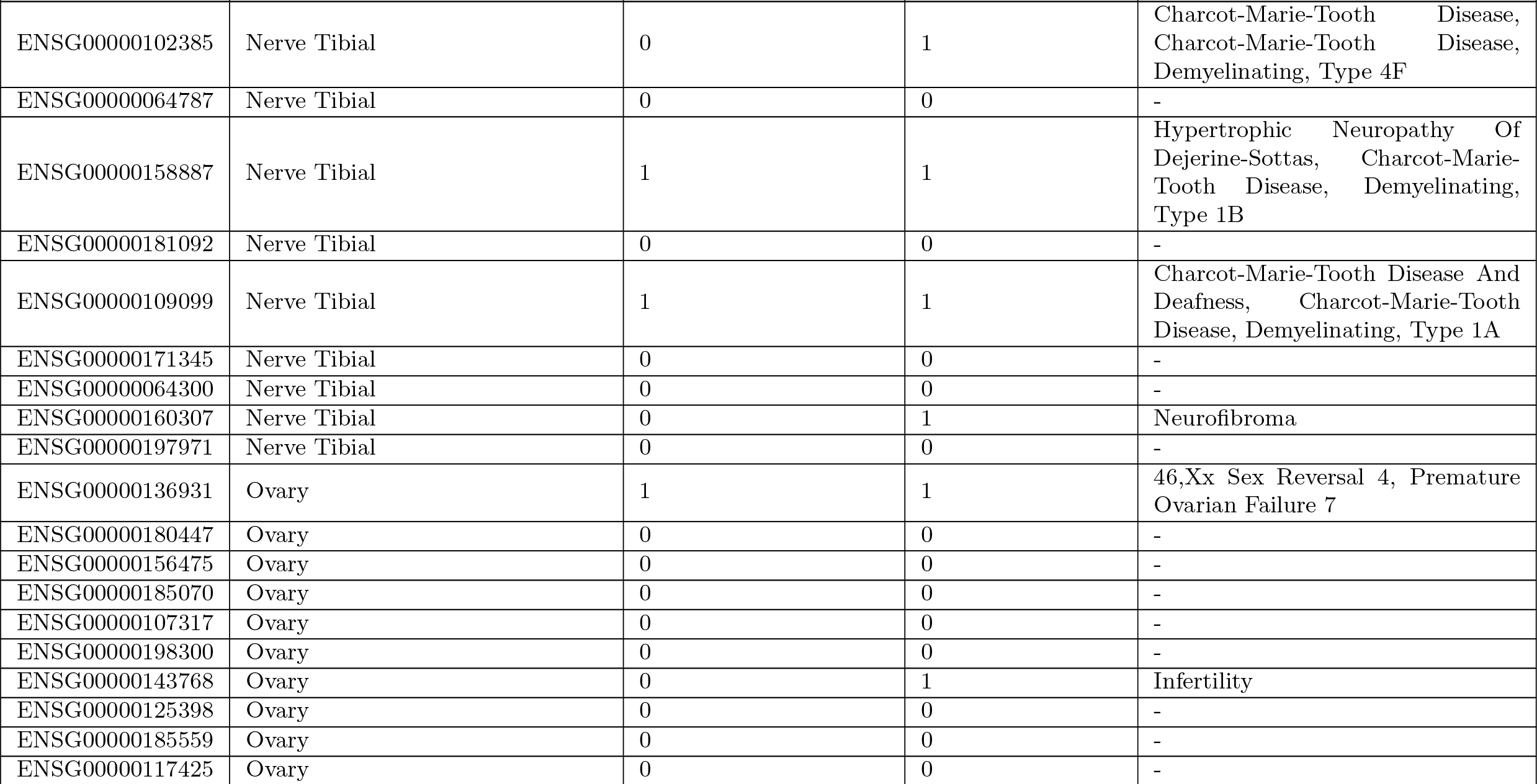
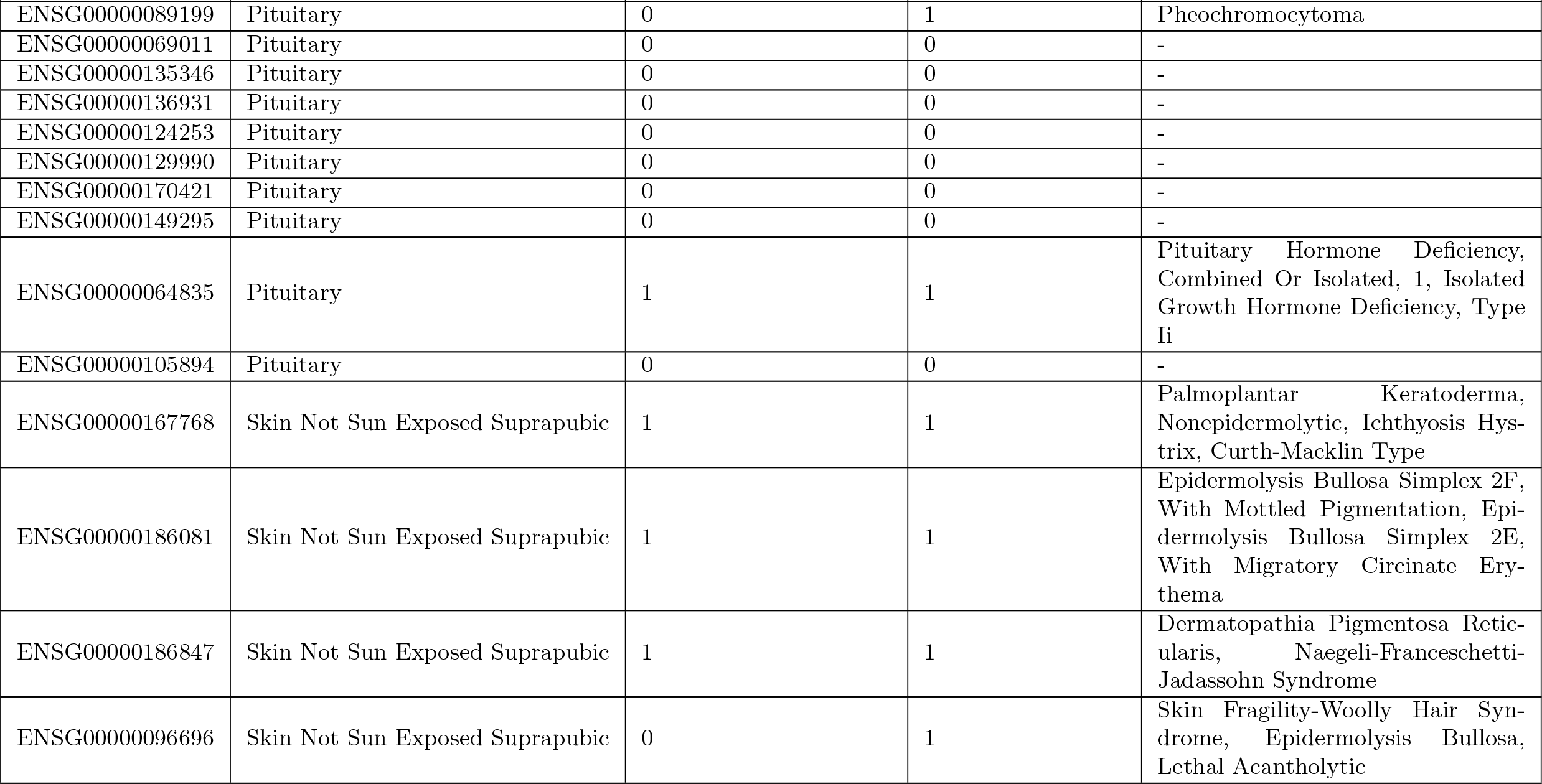
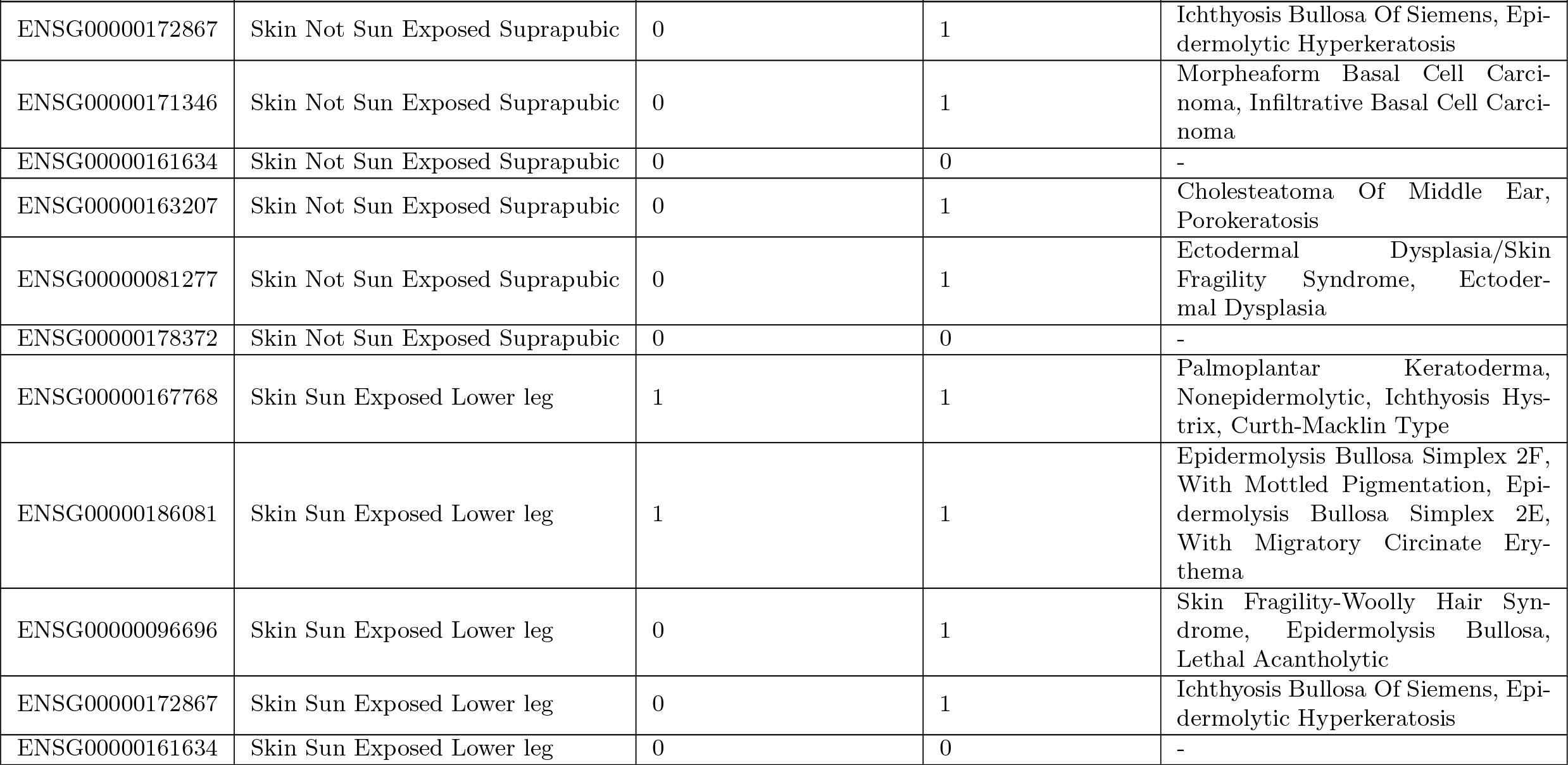
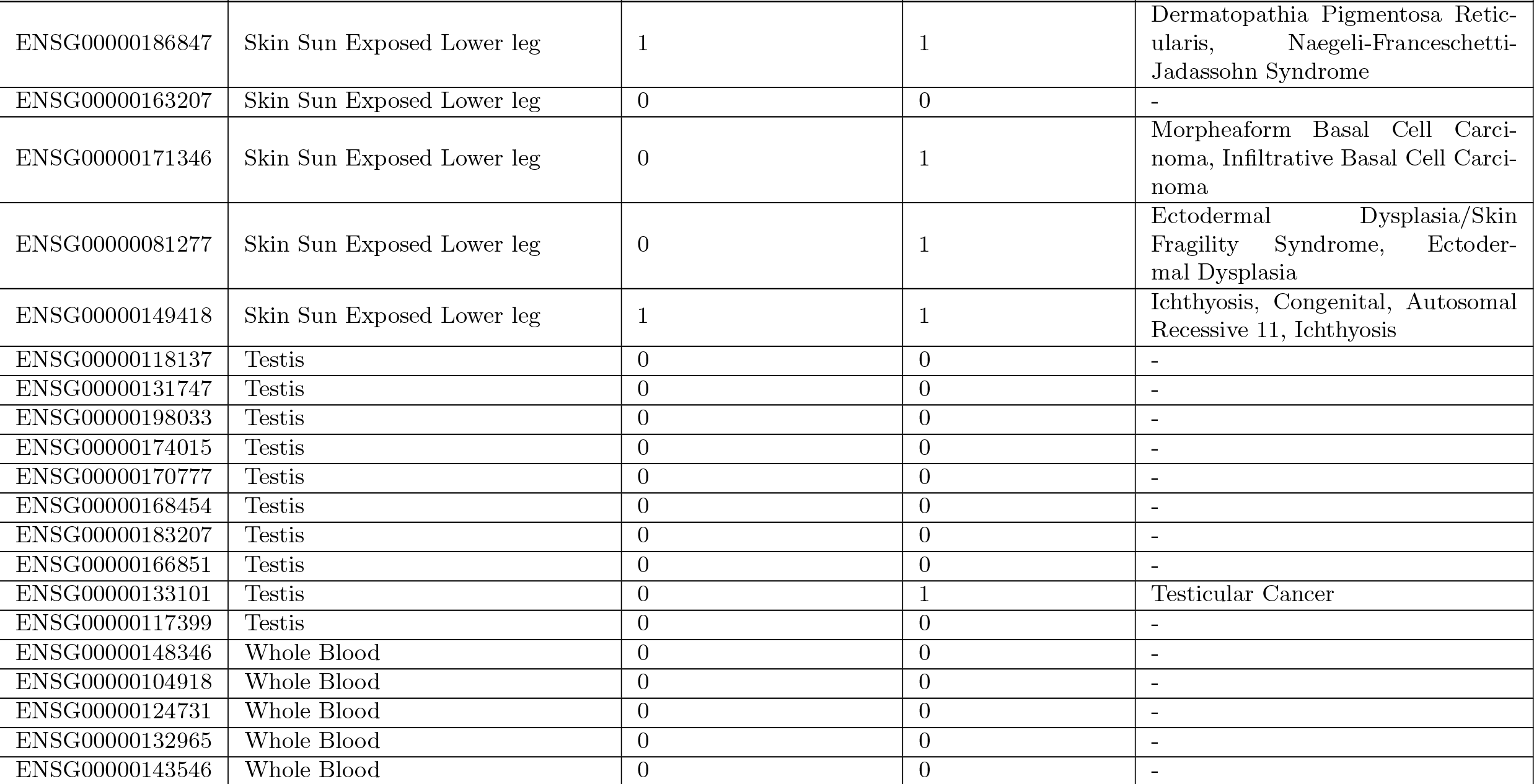
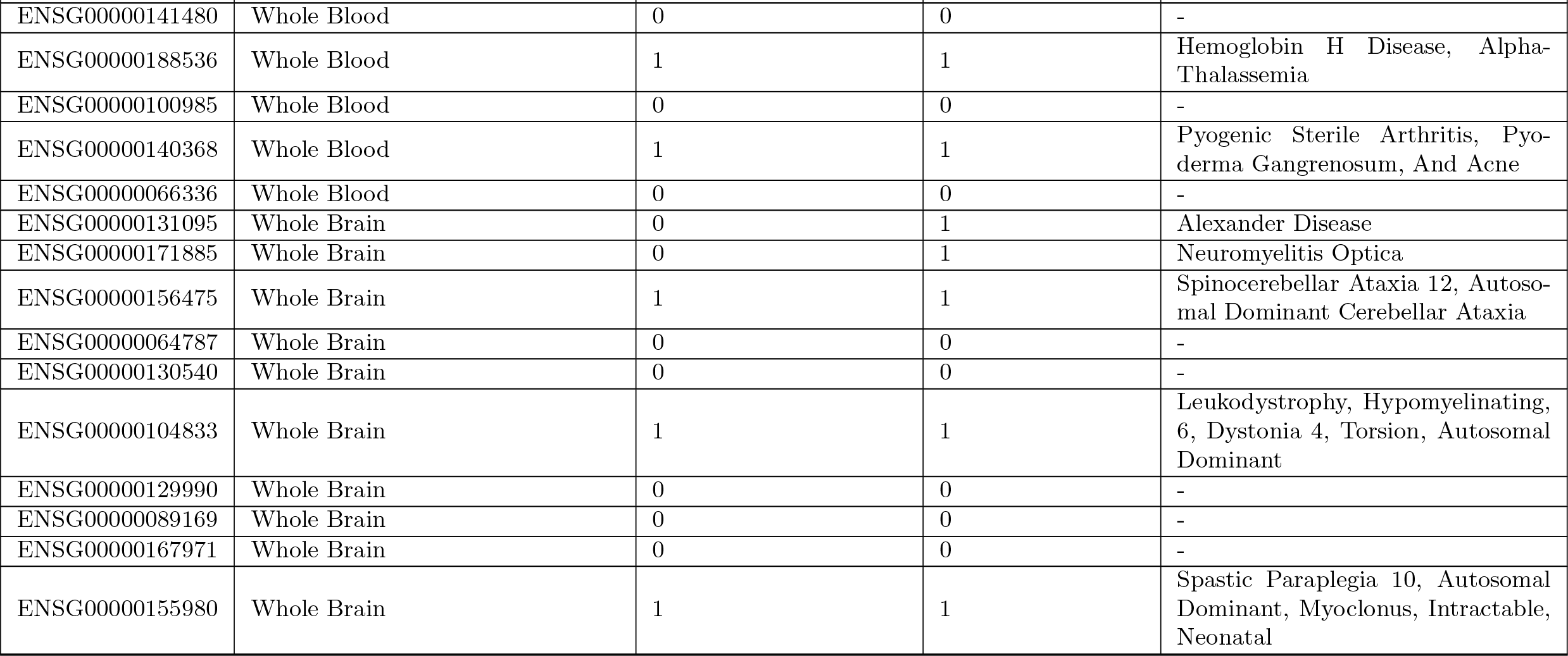
The top-ten genes predicted by the ensemble methods and the diseases that these genes were related to.

## Footnotes

1 Some node embedding algorithms, such as node2vec [15], use the edge weights when building the embedding. Using embedding created from a weighted graph to train an edge weight estimator creates a data leakage that will affect the whole system’s performance.

2 We tuned the parameters of the models by limiting the max depth of each tree and in this way, reducing overfit.

3 https://github.com/gloooryyt/oddball_py3

4 https://github.com/data4goodlab/wgand

